# Identifying Network Perturbation in Cancer

**DOI:** 10.1101/040394

**Authors:** Maxim Grechkin, Benjamin Logsdon, Andrew Gentles, Su-In Lee

## Abstract

We present a computational framework, called DISCERN (**DI**fferential **S**pars**E R**egulatory **N**etwork), to identify informative topological changes in gene-regulator dependence networks inferred on the basis of mRNA expression datasets within distinct biological states. DISCERN takes two expression datasets as input: an expression dataset of diseased tissues from patients with a disease of interest and another expression dataset from matching normal tissues. DISCERN estimates the extent to which each gene is *perturbed* – having distinct regulator connectivity in the inferred gene-regulatory dependencies between the disease and normal conditions. This approach has distinct advantages over existing methods. First, DISCERN infers *conditional dependencies* between candidate regulators and genes, where conditional dependence relationships discriminate the evidence for direct interactions from indirect interactions more precisely than pairwise correlation. Second, DISCERN uses a new likelihood-based scoring function to alleviate concerns about accuracy of the specific edges inferred in a particular network. DISCERN identifies perturbed genes more accurately in synthetic data than existing methods to identify perturbed genes between distinct states. In expression datasets from patients with acute myeloid leukemia (AML), breast cancer and lung cancer, genes with high DISCERN scores in each cancer are enriched for known tumor drivers, genes associated with the biological processes known to be important in the disease, and genes associated with patient prognosis, in the respective cancer. Finally, we show that DISCERN can uncover potential mechanisms underlying network perturbation by explaining observed epigenomic activity patterns in cancer and normal tissue types more accurately than alternative methods, based on the available epigenomic from the ENCODE project.

## Introduction

Genes do not act in isolation but instead work as part of complex networks to perform various cellular processes. Many human diseases including cancer are caused by dysregulated genes, with underlying DNA or epigenetic mutations within the gene region or its regulatory elements, leading to perturbation (topological changes) in the network [1–7]. This can ultimately impair normal cell physiology and cause disease [8–11]. For example, cancer driver mutations [12–19] on a transcription factor can alter its interactions with many of the target genes that are important in cell proliferation (Fig. 1A). A key tumor suppressor gene can be bound by different sets of transcription factors between cancer and normal cells, which leads to different roles [6,20–23] (Fig. 1B). Recent studies stress the importance of identifying the perturbed genes that create large topological changes in the gene network between disease and normal tissues as a way of discovering disease mechanisms and drug targets [8,9,24–27]. However, most existing analysis methods that compare expression datasets between different conditions (e.g., disease vs. normal tissues) focus on identifying the genes that are *differentially expressed* [28–30]. For example, a recent review paper on biological network inference [31] emphasized that there is a lack of methods that focus on inferring the differential network between different conditions (e.g., distinct species, and disease conditions).

**Figure 1.**
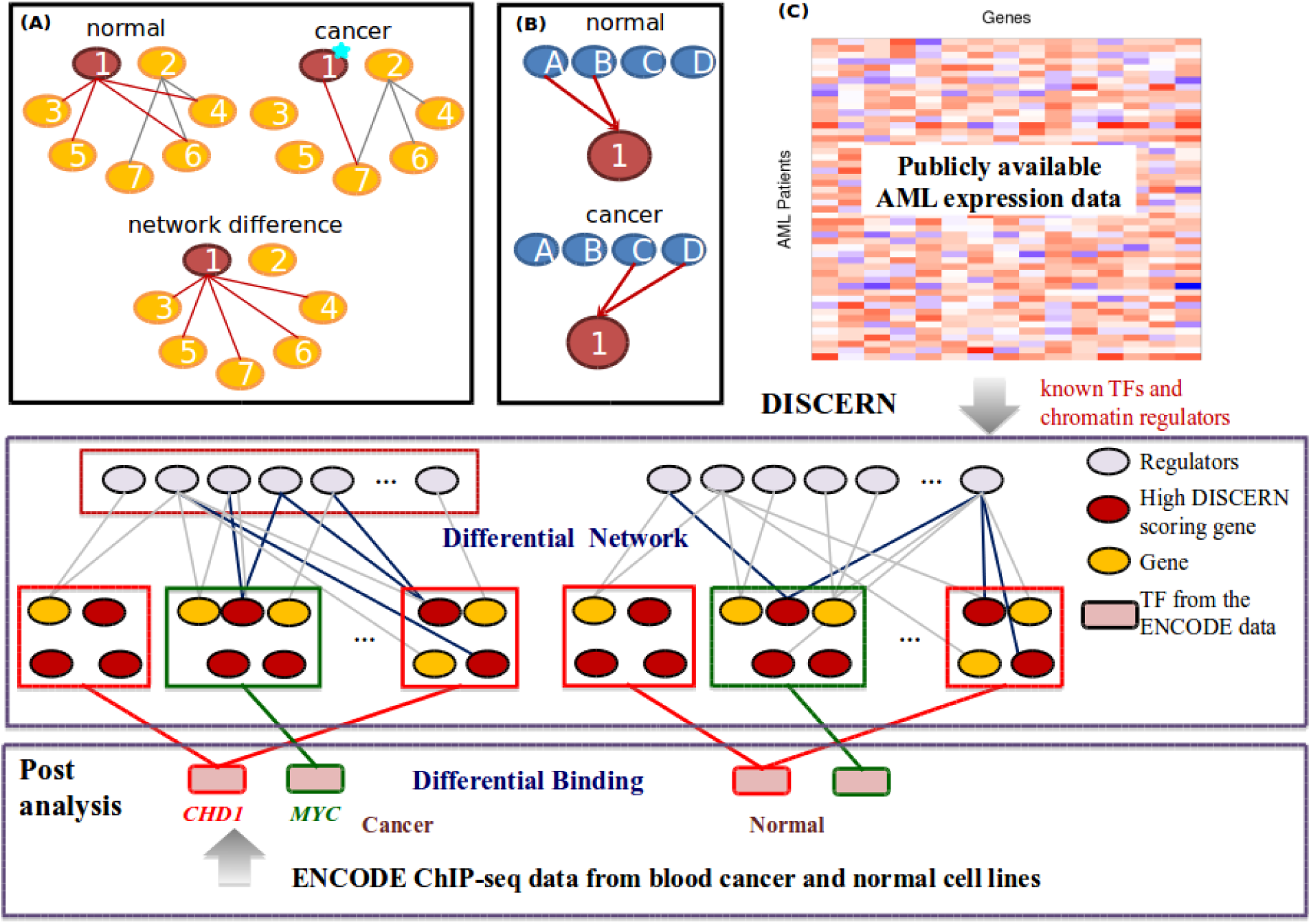
(A) A simple hypothetical example that represents the perturbation of the network of 7 genes between disease and normal tissues. One possible cause of network perturbation is a cancer driver mutation on gene ‘1’ that alters the interactions with genes ‘3’, ‘4’, ‘5’, and ‘6’. (B) One possible cause of network perturbation. Gene ‘1’ is regulated by different sets of genes between cancer and normal conditions. (C) The overview of our approach. DISCERN takes two expression datasets as input: an expression dataset from patients with a disease of interest and another expression dataset from normal tissues (top). DISCERN computes the score for each gene that estimates the difference in connection with other genes between disease and normal conditions (middle). We perform various post-analyses to evaluate the DISCERN method by comparing with alternative methods, based on the importance of the high-scoring genes in the disease through the survival analysis and on how well the identified perturbed genes explain the observed epigenomic activity data accurately (bottom).

Several recent studies compare gene networks inferred between conditions based on expression datasets [1, 32–37]. They fall into three categories: 1) Network construction based on prior knowledge: West et al. (2012) computes the local network entropy, based on the protein interaction network from prior knowledge and expression datasets from cancer and normal tissues [1]. 2) Pairwise correlation-based networks: Guan et al. (2013) [35] proposed the *local network similarity* (LNS) method that compares the pairwise correlation matrices of all genes between two conditions by using Pearson’s correlation. Various other authors compare pairwise correlation coefficients for all gene pairs between conditions with different methods, such as t-tests [32,33,36]. 3) Learning a condition-specific conditional dependency network and comparing between conditions: Gill et al. (2010) proposed a method, called PLSNet, that fits a partial least squares model to each gene, computes a connectivity scores between genes, and then calculates the *L*_1_ distance between score vectors to estimate network perturbation [34]. Zhang et al. (2009) proposed a differential dependency network (DDN) method that uses *lasso regression* to construct networks, followed by permutation tests [37]. There have been approaches to identify dysregulated genes in cancer by utilizing multiple types of molecular profiles, not based on network perturbation across disease states estimated based on expression data. Successful examples use a linear model to infer each gene expression model based on copy number variation, DNA methylation, ChIP-seq data, miRNAs or mRNA levels of transcription factors [38–40]. The advantages of the aforementioned methods that take only expression datasets as input to identify perturbed genes is in its applicability to data with many samples or to diseases for which only expression data are available. In this paper, we focus on identifying perturbed genes purely based on gene expression datasets representing distinct states, and compare our method with existing methods such as LNS, D-score and PLSNet.

We present a new computational method, called DISCERN (**DI**fferential **S**pars**E R**egulatory **N**etwork), to identify *perturbed* genes, i.e. the genes with differential connectivity between the condition specific networks (e.g., disease versus normal). DISCERN takes two expression datasets, each from a distinct condition, as input - where it computes a novel perturbation score for each gene. The *perturbation score* captures how likely a given gene is to have a distinct set of regulators between conditions (Fig. 1A). The DISCERN method contains specific features that provide advantages over existing approaches: 1) DISCERN can distinguish direct associations among genes from indirect associations more accurately than marginal association methods such as LNS; 2) DISCERN uses a penalized regression-based modeling strategy that allows efficient inference of genome-wide gene regulatory networks; and 3) DISCERN uses a new likelihood-based score that is more robust to the expected inaccuracies in local network structure estimation. We elaborate on these three advantages below:

First, DISCERN infers gene networks based on *conditional dependencies* among genes - a key type of probabilistic relationship among genes that is fundamentally distinct from correlation. If two genes are conditionally dependent, then by definition, their expression levels are still correlated even after accounting for (e.g., regressing out) the expression levels of all other genes. Thus, conditional dependence relationship is less likely to reflect transitive effects than mutual correlation, and provides stronger evidence that those genes are functionally related. These functional relationships could be regulatory, physical, or other molecular functionality that causes two genes expression to be tightly coupled. As a motivating example, assume that the expression levels of genes ‘3’ and ‘5’ are regulated by gene ‘1’ in a simple 7-gene network (Fig. 1A). This implies that the expression level of gene ‘1’ contains sufficient information to know the expression levels of genes ‘3’ and ‘5’. In other words, genes ‘3’ and ‘5’ are *conditionally independent* from each other and from the rest of the network given gene ‘1’.

Second, DISCERN uses an efficient neighborhood selection strategy based on a penalized regression to afford inference of genome-wide networks. Penalized regression is a well established technique to identify conditional dependencies [41]. Inferring the conditional dependence relationships from high-dimensional expression data (i.e., where the number of genes is much greater than the number of samples) is a challenging statistical problem, due to a very large number of possible network structures among tens of thousands of genes. Unlike pairwise correlation, the conditional dependence between ‘1’ and ‘2’ cannot be measured based on just the expression levels of these two genes. One must consider a number of possible networks among all genes and find the one that best explains the expression data. This involves both computational and statistical challenges. To make this process feasible, DISCERN uses a sparse regression model for each gene to select neighbors in the network [41,42]. The inference of the genome-wide network with a conditional dependence method is a key distinguishing feature of the DISCERN method.

Finally, one of the most novel features of DISCERN is a network perturbation score that avoids overestimation of the degree of network perturbation due to dense correlation among many genes. Revisiting the 7-gene network (Fig. 1A), assume that genes ‘5’ and ‘7’ are highly correlated to each other, in which case a penalized regression that imposes a sparsity prior, such as the *lasso* method, may arbitrarily select one of them. This can result in a false positive edge between genes ‘1’ and ‘7’ instead of ‘1’ and ‘5’. This may lead to overestimation of the perturbation of gene ‘1’ (Fig. 1A). Our network perturbation score overcomes this limitation by measuring the network differences between conditions based on the likelihood when the estimated networks are swapped between conditions - not based on the differences in topologies of the estimated networks. We demonstrate the effectiveness of this feature by comparing with methods based on the topology differences of the estimated networks.

We evaluate DISCERN on both synthetic and gene expression data from three human cancers, acute myeloid leukemia (AML), breast cancer (BRC), and lung cancer (LUAD). Integrative analysis using DISCERN on epigenomic data from the Encyclopedia of DNA Elements (ENCODE) project leads to hypotheses on the mechanisms underlying network perturbation (Fig. 1C). The resulting DISCERN score for each gene in AML, BRC and LUAD, the implementation of DISCERN, and the data used in the study are freely available on our website discern-leelab.cs.washington.edu/.

## Results

### Method overview

Here, we describe the DISCERN method, referring to the Methods for a full description. We postulate that a gene can be perturbed in a network largely in two ways: A gene can change how it influences other genes (Fig. 1A), for example transcription factor mutations affecting cell proliferation pathways and possibly causing cancer [12–19]. A gene can change the way it is influenced by other genes, a common example being when a mutated (genetically or epigenetically) gene acquires a new set of regulators, which occurs frequently in development and cancer [6,20–23] (Fig. 1B). Identifying the genes that are responsible for large topological changes in gene networks when perturbed could be crucial for understanding disease mechanisms and identifying key drug targets [8,9,24–27]. However, most current methods for identifying these genes rely on *differential expression* [28–30], rather than genes being *differently connected (rewired)* to other genes in the gene expression network (Fig. 1A).

We model all genes’ expression values using sparse linear models (*lasso* regressions): let 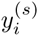 be expression levels of gene *i* in an individual with state *s*, cancer (*s* = *c*) or normal (*s* = *n*), modeled as: 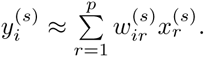.

Here, *x*_1_,…*x*_p_ denote *candidate regulators*, a set of genes known to regulate other genes, including transcription factors, chromatin modifiers or regulators, and signal transduction genes, which were used in previous work on network reconstruction approaches [43–46] **(S1 Table).** Linear modeling allows us to capture conditional dependencies efficiently from genome-wide expression data containing tens of thousands genes. Naturally, a zero weight *wi*_r_ indicates that a regulator *r* does not affect the expression of the target gene *i.* Sparsity-inducing regularization helps to select a subset of candidate regulators, which is a more biologically plausible model than having all regulators, and makes the problem well-posed in our *high-dimensional* setting (i.e., number of genes ≫ number of samples).

To determine the regulators for any given gene, we use a lasso penalized regression model [47] with the optimization problem for each lasso regres sion defined as 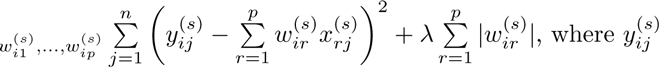 corresponds to the expression level of the *i*^th^ gene in the *j*^th^ patient in the *s*^th^ state, and 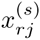 similarly corresponds to the expression level of the *r*^th^ regulator in the *j*^th^ patient in the *s*^th^ state. The second term, the *L*_1_ penalty function, will zero out many irrelevant regulators for a given gene, because it is known to induce sparsity in solution [47]. We normalize the expression levels of each gene and each regulator to be mean zero and unit standard deviation, a technique called standardization, which is a standard practice before applying a penalized regression method [47–50]. The difference in the weight vector between conditions, 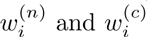, could indicate distinct connectivity of gene i with p regulators between the conditions. However, simply computing the difference of the weight vectors is unlikely to be successful, due to the correlation among regulators. The lasso, or other sparsity-inducing regression methods, can arbitrarily choose different regulators between cancer and normal. Examining the difference in the weight vectors between conditions would therefore lead to overestimation of network perturbation.

Instead, DISCERN adopts a novel network perturbation score that measures how well each weight vector learned in one condition explains the data in a different condition. This increases the robustness of the score to correlation among regulators, as demonstrated in the next section. We call this score the DISCERN score, defined as 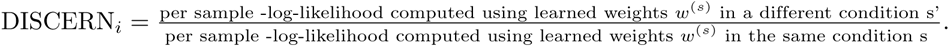. This is equivalent to 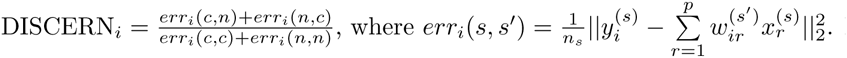. Here *n*_s_ is the number of samples in the data from condition *s*. The numerator measures the error of predicting gene *i*’s expression levels in cancer (normal) based on the weights learned in normal (cancer). If gene *i* has different sets of regulators between cancer and normal, it is likely to have a high DISCERN score. The denominator plays an important role as a normalization factor, which is demonstrated by comparing with an alternative score, namely the D^0^ score (Fig. 2A), that uses only the numerator of the DISCERN score. We also compare with existing methods, such as LNS [35] and PLSNet [34], that compare the weight vectors between cancer and normal models where we demonstrate the advantages of the likelihood-based model that DISCERN uses.

**Figure 2.**
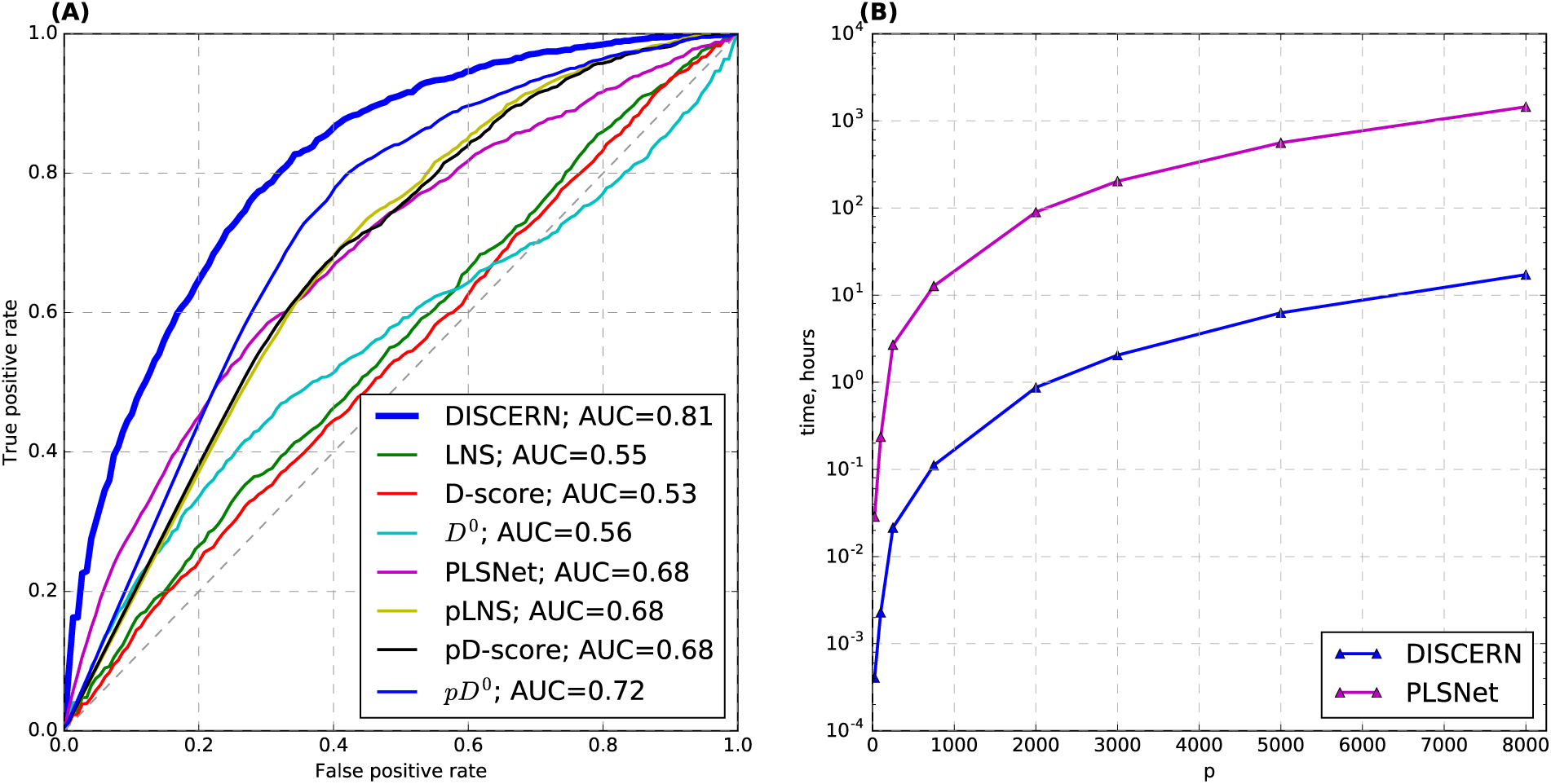
(A) Average receiver operating characteristic (ROC) curves from synthetic data experiments. We compare DISCERN with 7 alternative methods: 3 existing methods – LNS [35], D-score [36], and PLSNet [34] – and 4 methods we developed for comparison – pLNS, pD-score, *D*^0^ and p*D*^0^. (B) Comparison of the runtime (hours) between PLSNet and DISCERN over varying numbers of variables. The triangles mean the measured run times over specific values of *p*, the number of variables, and lines connect these measured run times. PLSNet uses the empirical p-values from permutation tests as scores, and DISCERN does not. We show that DISCERN is two to three orders of magnitude faster than PLSNet.

### Comparison with previous approaches on synthetically generated data

In order to systematically compare DISCERN with alternative methods in a controlled setting, we performed validation experiments on 100 pairs of synthetically generated datasets representing two distinct conditions. Each pair of datasets contains 100 variables drawn from the multivariate normal distribution with zero mean and covariance matrices Σ_1_ and Σ_2_. We divided 100 variables into the following three categories: 1) variables that have different sets of edge weights with other variables across two conditions, 2) variables that have exactly the same sets of edge weights with each other across the conditions, and 3) variables not connected with any other variables in the categories 2) and 3) in both conditions. For example, in Fig. 1A, ‘1’ is in category 1). ‘2’, ‘4’, ‘6’, and ‘7’ are in category 2), and ‘3’ and ‘5’ is in category 3). We describe how we generated the network edge weights (i.e., elements of 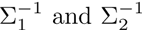) among the 100 variables in more detail in Methods.

We compared DISCERN with 4 alternative methods to identify perturbed genes: LNS [35], D-score [36], PLSNet [34], and *D*^0^ that uses only the numerator of the DISCERN score. Here, we do not compare with the methods to identify differentially expressed genes, such as ANOVA, because the synthetic data were generated from a zero mean Gaussian distribution. We note that the PLSNet method uses empirical p-values as the network perturbation scores, where the empirical p-value for each gene is estimated from permutation tests that generate the null distribution of the gene’s score [34]. All the other methods, such as DISCERN, LNS, and D-score, do not require permutation tests (see Methods for details). To show that DISCERN outperforms existing methods and those that use the empirical p-values obtained through permutation tests as the network perturbation scores, we developed the following methods for comparison: LNS (D-score, or *D*^0^) followed by permutation tests to compute the empirical p-values, called pLNS, pD-score, and p*D*^0^, respectively.

The averaged receiver operating characteristic (ROC) curves across 100 pairs of datasets for these methods (Fig. 2A) show that DISCERN significantly outperforms all the other 7 methods – 3 existing methods (LNS, D-score, and PLSNet), and 4 methods we created for comparison (*D*^0^, p*D*^0^, pLNS, and pD-score). Except DISCERN, PLSNet performs the best among all existing methods. However, its run time grows too quickly as the number of variables increases, which makes it two to three orders of magnitude slower than DISCERN when run on larger data (Fig. 2B). PLSNet was too slow to run on genome-scale data and therefore we did not use it for the subsequent experiments on genome-wide gene expression data from cancer patients.

We note that DISCERN does not need permutation tests to generate the null distribution of the score for each gene. All other methods improve when the empirical p-values from permutation tests are used, which indicates that the gene-level bias on the magnitude of the raw scores hurts their performance to identify perturbed genes. DISCERN significantly outperforms *D*^0^ that uses only the numerator of the DISCERN score, which indicates that the denominator of the DISCERN score plays a role to normalize the score such that the scores of different genes can be comparable. Computing the empirical p-value for each gene based on the gene-specific null distribution obtained through permutation tests is not feasible on genome-wide data. To obtain a p-value of 0.05 after Bonferroni correction, we need at least (1/0.05 × *p*) permutation tests per gene, where *p* is the total number of genes, and (1/0.05 × *p*^2^) permutation tests in total. When *p* = 20, 000, this number is (4 × 10^9^) permutation tests, which is not feasible even when using multiple processors at a reasonable cost. This is demonstrated in Fig. 2B that shows the run time of PLSNet, a permutation test-based method, when applied to data containing a varying number of genes (*p*).

### Comparison of methods on gene expression datasets

We used genome-wide expression datasets consisting of 3 acute myeloid leukemia (AML) datasets, 3 breast carcinoma (BRC) datasets and 1 lung adenocarcinoma (LUAD) dataset (Table 1). Details on the data processing are provided in Methods. To evaluate the performance of the DISCERN method, we compared with existing methods that scale to over tens of thousands of genes: LNS [35] and D-score [36] that aim to estimate network perturbation, and ANOVA that measures differential expression levels between cancer and normal samples.

**Table 1.** Gene expression datasets used in this paper.

|  | Reference | # Genes | # of Tumors | # of Normal | Survival? | Platform | Accession |
| --- | --- | --- | --- | --- | --- | --- | --- |
| AML1 | MILE, [53] | 16853 | 541 | 73 | No | Affy U133+2.0 | GSE13159 |
| AML2 | Gentles, [54] | 16853 | 515 | 0 | Yes | Affy U133+2.0 | GSE12417,GSE14468,GSE1 |
| AML3 | Metzeler | 11697 | 152 | 0 | Yes | Affy U133A | GSE12417 |
| BRC1 | TCGA | 10809 | 529 | 61 | Yes | Agilent G4502A | Firehose 2013042100 |
| BRC2 | Metabric | 10809 | 1981 | 0 | Yes | Illumina HTv3 | Synapse: syn1688369 |
| BRC3 | Oslo | 10809 | 184 | 0 | Yes | Illumina HTv3 | Synapse: syn1688370 |
| LUAD1 | TCGA | 17022 | 504 | 57 | Yes | Illumina HiSeq | Firehose 2015110100 |

We first computed the DISCERN, LNS, D-score, and ANOVA scores in the 3 cancers based on the following datasets that contain normal samples: AML1, LUAD1 and BRC1. Then, we used the rest of the datasets to evaluate the performance of each method at identifying genes previously known to be important in the disease, for example, the genes whose expression levels are significantly associated with survival time in cancer. The value of the sparsity tuning parameter λ was chosen via cross-validation tests, a standard statistical technical to determine the value of λ [47].

To remove any potential concern of the effect of standardization on genes with very low expression level, we first show that genes with low mean expression do not tend to have high enough DISCERN score to be considered in our evaluation in the next sections (S1 Fig). The Pearson’s correlation between the mean expression before standardization and the DISCERN score ranges from 0.08 and 0.43. Positive correlation is induced because genes with low mean expression tend to have lower DISCERN scores, indicating that there is probably not an issue in terms of overly selecting genes whose expression are likely essentially noise. To further reduce the potential concern of genes with low expression in RNA-seq data (LUAD), we applied the voom normalization method that is specifically designed to adjust for the poor estimate of variance in count data, especially for genes with low counts [51].

We assessed the significance of the DISCERN scores through a conservative permutation testing procedure, where we combined cancer and normal samples, and permuted the cancer/normal labels among all samples (more details in Methods). Unlike the gene-based permutation test described in the previous section, here, we generated a single null distribution for all genes, which requires a significantly less number of permutation tests (one million in this experiment). After applying false discovery rate (FDR) correction on these p-values, there are 1,351 genes (AML), 2,137 genes (BRC), and 3,836 (LUAD) genes whose FDR corrected p-values are less than 0.05. We consider these genes to be significantly perturbed genes (S2 Table). The difference in these numbers of significant perturbed genes identified by DISCERN is consistent with a prior study that showed that lung cancer has a larger number of non-synonymous mutations per tumor than breast cancer, which has a larger number than AML [52].

### Top scoring DISCERN genes in AML reveal known cancer drivers in AML

The 1,351 genes that were predicted to be significantly perturbed between AML samples and normal non-leukemic bone marrow samples were enriched for genes causally implicated previously in AML pathogenesis (S2 Table). We noted deregulation of a number of genes that we and others have previously identified as being aberrantly activated in leukemic stem cells including BAALC, GUCY1A3, RBPMS, and MSI2 [55–57]. This is consistent with over-production of immature stem-cell like cells in AML, which is a major driver of poor prognosis in the disease. Prominent among high-scoring DISCERN genes were many HOX family members, which play key roles in hematopoietic differentiation and in the pathogenesis of AML [58]. HOX genes are frequently deregulated by over-expression in AML, often through translocations that result in gene fusions. The highest ranked gene in AML by DISCERN is HOXB3 which is highly expressed in multipotent hematopoietic progenitor cells for example. Thirteen (out of 39 known) HOX genes are in the 1,351 significantly perturbed genes (p-value: 5.99 × 10^−6^).

When compared to known gene sets from the Molecular Signature Database (MSigDB) [59] in an unbiased way, the top hit was for a set of genes (VERHAAK AML WITH NPM1 MUTATED DN; p-value: 2 × 10^−86^) that are down-regulated by NPM1 (nuclephosmin 1) mutation in AML (see Supporting Information). NPM1 is one of three markers used in AML clinical assessment; the others are FLT3 and CEBPA that are significantly perturbed genes identified by DISCERN as well. Mutation leads to aberrant cytoplasmic location of itself and its interaction partners, leading to changes in downstream transcriptional programs that are being captured by DISCERN. Also highly significant were genes highly expressed in hematopoietic stem cells [60] (JAATINEN HEMATOPOIETIC STEM CELL UP; p-value: 6 × 10^−74^). Among these were key regulators of hematopoietic system development such as KIT, HOXA3, HOXA9, HOXB3 (with the latter homeobox genes also implicated in AML etiology); as well as FLT3 which plays a major role in AML disease biology, with its mutation and constitutive activation conferring significantly worse outcomes for patients [61]. Comparison to Gene Ontology (GO) categories identified dysregulation of genes involved in hemostasis and blood coagulation, a key clinical presentations of AML. Furthermore, GTPase activity/binding and SH3/SH2 adaptor activity were enriched among DISCERN genes (see Supporting Information). These are pertinent to AML due to previously noted high expression in AML leukemic stem cells of GUCY1A3 and SH3BP2, both identified as perturbed genes by DISCERN [55]. However, their function has not been examined in detail, suggesting that they are potential targets for further investigation as to their role in AML disease mechanisms. Several other highly significant enrichments were for AML subtypes that are driven by specific translocations, including MLL (mixed lineage leukemia) translocation with various partners, as well as t(8;21) translocations. The latter is of particular interest, since it is primarily a pediatric AML, whereas our network analysis uses purely adult AML samples – indicating the potential to uncover putative mechanisms that generalize beyond the context of the immediate disease type.

### Top scoring DISCERN genes in lung cancer reveal biological processes known to be important in lung cancer

There are 3,836 significantly perturbed genes identified by DISCERN in lung cancer (LUAD) (S2 Table). The 3rd and 4th highest ranked genes in LUAD are ICOS (inducible costimulator), YWHAZ (14-3-3-zeta). Both have described roles in disease initiation or progression in lung cancer. Polymorphisms in ICOS have been associated with pre-disposition to non-small cell lung cancer [62]; while over-expression of YWHAZ is known to enhance proliferation and migration of lung cancer cells through induction of epithelial-mesenchymal transitions via beta-catenin signaling [63]. GIMAP5 (GTPase IMAP Family Member 5), another high scoring LUAD gene (11th), is consistently repressed in paired analyses of tumor vs normal lung tissue from the same patient, and encodes an anti-apoptotic protein [64]. Down-regulation of GIMAP5 in lung tumors therefore potentially facilitates their evasion of programmed cell death, one of the hallmarks of cancer.

Several of the GO biological categories enriched in 3,836 high-scoring DISCERN genes in LUAD (FDR-corrected p-value < 0.05) reflected metabolic and proliferative processes that are commonly de-regulated in solid tumors such as lung adenocarcinoma. Among these were cellular response to stress, mitotic cell cycle, amino acid metabolism, and apoptosis (see Supporting Information). In fact the top-ranked gene was MCM7 (minichromosome maintenance protein 7), an ATP-dependent DNA helicase involved in DNA replication which has been implicated in carcinogenesis previously due to its function as a binding partner of PRMT6 [65]. Moreover, it was specifically identified as being a potential therapeutic target due to its over-expression in solid tumors relative to normal tissues. The high ranking of genes associated with apoptosis is consistent with the fact that there is often high rate of tumor cell death. Although the highly-ranked CARD6 (caspase recruitment domain family member 6) functions in apoptotic processes, it is also known as a regulator of downstream NF-*κβ* signaling. Indeed, consistent with this, we found enrichment for NF-*κβ* signaling pathway genes among high DISCERN-scoring genes in LUAD including NFKBIB (NF-*κβ* inhibitor *β*) which inhibits the NF-*κβ* complex by “trapping” it in the cytoplasm, preventing nuclear activation of its downstream targets. Although the role of NFKBIB in lung cancer has not been studied extensively, its related family member NFKBIA is known to be a silencer in non-small-cell lung cancer patients with no smoking history, suggesting that it could play some role in LUAD that arises through inherent genetic influences, or environmental insults other than smoking [66]. Levels of *β*-catenin have been known for some time to influence progression and poor prognosis in LUAD, potentially through its role in differentiation and metastasis from primary tumor sites [67]. We found that components of *β*-catenin degradation pathways - including most notably CTNNBIP1 (β-catenin interacting protein 1) - ranked among the most significant DISCERN genes in our LUAD analysis.

When comparing to other sets of genes in MSigDB, we also found targets of transcription factors including MYC, which is often de-regulated in solid tumors (either by mutation or copy number variation), and targets of the polycomb repressive complex gene EZH2. The developmental regulator EZH2 functions through regulation of DNA methylation [68], and has been implicated in B-cell lymphomas through somatic mutations [69], promotion of transformation in breast cancer [70], as well as progression in prostate cancer [71]. Interestingly, the most highly dys-regulated gene set identified by comparison to GO categories in LUAD was one related to NGF (nerve growth factor)-TrkA signaling. There are a few reports of the relevance of this axis to cancers including neuroblastoma, ovarian cancer, and a possible role in promoting metastasis in breast cancer. However, its striking appearance as the most significant hit for high-ranking DISCERN genes suggests that it merits study in LUAD.

### Top scoring DISCERN genes in breast cancer reveal biological processes known to be important in breast cancer

Here, we did the functional enrichment analysis with 2,137 genes identified by DISCERN to be significantly perturbed in breast cancer (BRC) (S2 Table). BRC showed perturbation of distinct genes and sets of genes in comparison to LUAD, as well as similarities. Again, these included GO biological processes that one would generically expect to be over-activated in a solid tumor, such as translation intiation, cell cycle, proliferation, and general cellular metabolic processes. As with LUAD, targets of MYC were enriched in high-scoring DISCERN genes. Another high-scoring group in BRC was comprised of genes that are highly correlated with each other, but with this relationship de-regulated by BRCA1 mutation [72]. Additional significant overlaps were identified with luminal A, luminal B, HER2-enriched, and basal-like breast cancer subtype-specific genes that are associated with clinical outcomes [73]; and with genes associated with ER-positive breast cancer [74]. The 3rd highest ranked DISCERN gene was BRF2 (TFIIB-related factor 2). BRF2 is a known oncogene in both breast cancer and lung squamous cell carcinoma, and a core RNA polymerase III transcription factor that senses and reacts to cellular oxidative stress [75]. A GO category associated with NGF (nerve growth factor)-TrkA signaling shows the highest overlap with DISCERN genes in BRC (p-value: 3.16 × 10^−104^). NGF-TrkA signaling is upstream of the canonical phosphatidylinositol 3-kinase (PI3K)–AKT and RAS–mitogen-activated protein kinase (MAPK) pathways, both of which impinge on cell survival and differentiation. In the context of breast cancer, over-expression of TrkA has been connected to promoting growth and metastasis, as an autocrine factor, presumably due to its influence on PI3K-AKT and RAS/MAPK [76]. TrkA is reportedly over-expressed in breast carcinoma relative to normal breast tissue in a majority of cases [77], supporting the high-ranking of genes in this pathway by DISCERN. Taken together, these results indicate that DISCERN highly ranks genes that are connected to known phenotypic and survival-associated processes in breast cancer. However, intriguingly the top DISCERN gene was CLNS1A (chloride nucleotide-sensitive channel 1A). This chloride channel gene has not, to our knowledge, been implicated in pathogenesis in any cancer, although it is a member of the BRCA1-related correlation network noted above. In fact there appear to have been few studies of its function although Entrez gene notes that it “performs diverse functions”.

### DISCERN scores reveal survival-associated genes across multiple cancer types

In this section, we focus on the quantitative assessment of DISCERN and the comparison with LNS and D-score in terms of how much the identified genes are enriched for genes implicated to be important in the disease. Specifically, genes whose expression levels are significantly associated positively or negatively with survival time are often considered to be associated with tumor aggression. Identifying such genes has been considered as an important problem by a number of authors, where breast cancer was one of the first cancers to show promise in terms of identifying clinically relevant biomarkers [78,79]. Here, we evaluated DISCERN based on how well it reveals survival-associated genes identified in an available independent dataset.

We chose the datasets with measures of patient prognosis: AML2, BRC2, and LUAD1. AML2 and BRC2 were not used for computing any scores (DISCERN, LNS, D-score, and ANOVA). For each of these datasets we computed the survival p-values based on the Cox proportional hazards model [80] measuring the association between each gene’s expression level and survival time. We defined survival-associated genes as the genes whose expression levels are associated with survival time based on the Cox proportional hazards model (p-value < 0.01) (**S3 Table**).

We considered the genes whose DISCERN scores are significantly high at FDR corrected p-value < 0.05 in each cancer: 1,351 genes (AML), 2,137 genes (BRC), and 3,836 genes (LUAD). We first computed the Fisher’s exact test p-values to measure the statistical significance of the overlap between these significantly perturbed genes and survival-associated genes in each of three cancers. For each cancer, we compared with existing methods to detect network perturbation – LNS and D-score – when exactly the same number of top-scoring genes were considered (Fig. 3A-C). Since these numbers of genes were chosen specifically for DISCERN, there is a chance that LNS and D-score would show a higher enrichment for survival-associated genes if different numbers of top-scoring genes were considered. As discussed in the previous section, performing the gene-based permutation tests to estimate the confidence of each gene’s score in genome-wide data is not feasible. Instead, we compared the Fisher’s exact test p-values of the three methods across a range of numbers of top-scoring genes from 0 to N Fig. 3D-F. It is pretty clear that neither LNS nor D-score would be better than DISCERN in revealing survival-associated genes, even when different numbers of top-scoring genes were considered across all cancer types.

**Figure 3.**
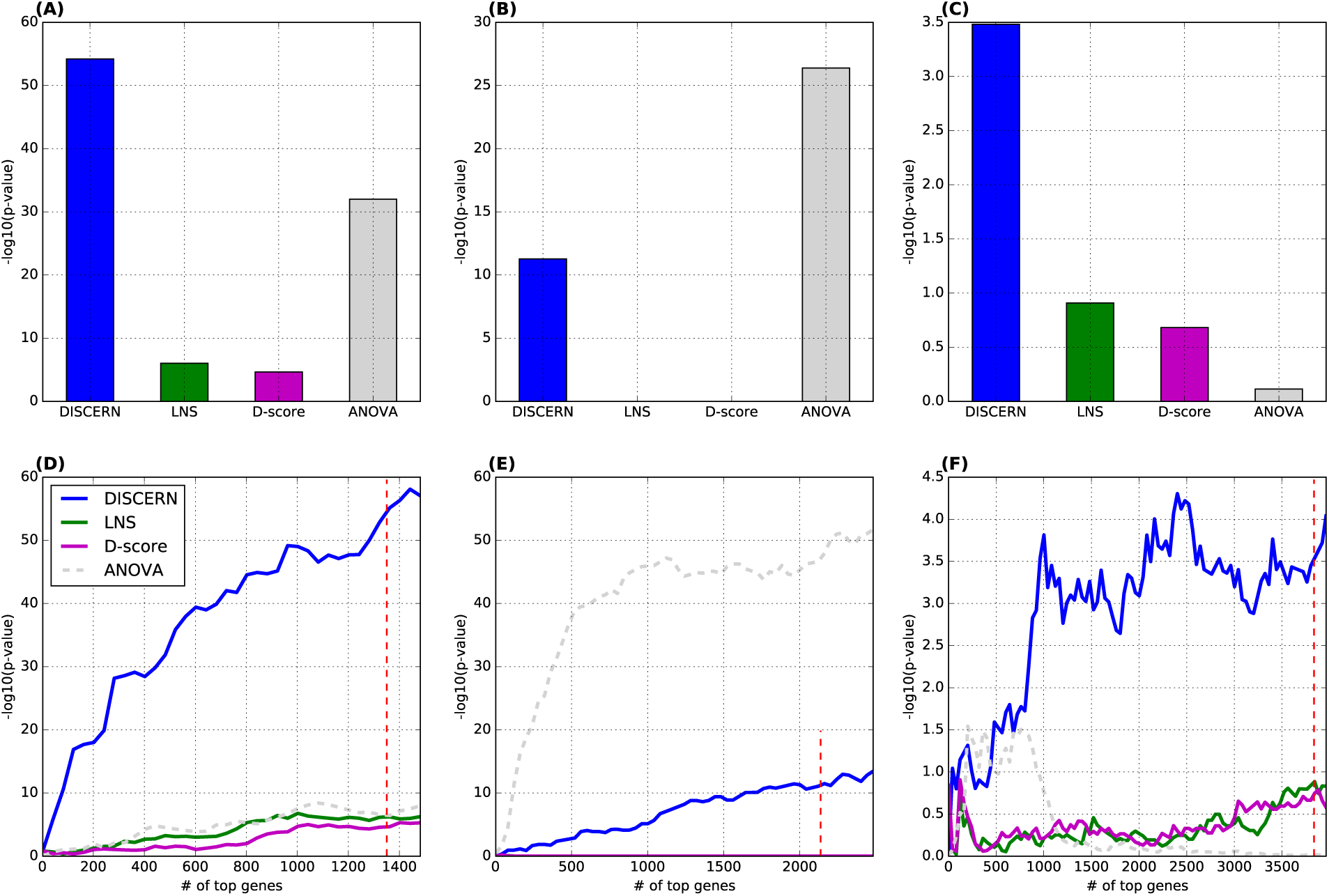
The significance of the enrichment for survival-associated genes in the identified perturbed genes. We compared DISCERN with LNS and D-score based on the Fisher’s exact test p-value that measures the significance of the overlap between N top-scoring genes and survival-associated genes in each of three cancers. (A)-(C) We plotted-log_10_(p-value) from the Fisher’s exact test when N top-scoring genes were considered by each method in 3 datasets: (A) AML (N = 1,351), (B) BRC (N = 2,137), and (C) LUAD (N = 3,836). For ANOVA, we considered 8,993 genes (AML), 7,922 genes (BRC) and 13,344 genes (LUAD) that show significant differential expression at FDR corrected p-value < 0.05. (D)-(F) We consider up to 1,500 (AML), 2,500 (BRC), and 4,000 (LUAD) top-scoring genes in each method, to show that DISCERN is better than LNS and D-score in a range of N value. The red-colored dotted line indicates 1,351 genes (AML), 2,137 genes (BRC), and 3,836 genes (LUAD) that are identified to be significantly perturbed by DISCERN (FDR < 0.01). We compare among the 4 methods consisting of 3 methods to identify network perturbed genes (solid lines) and ANOVA for identifying differentially expressed genes (dotted line) in 3 cancer types.

ANOVA is a well-established method to identify *differentially expressed* genes across distinct conditions; DISCERN LNS, and D-score are methods to identify *differentially connected* genes across conditions. Therefore, the purpose of the comparison with ANOVA is not to evaluate DISCERN in identifying survival-associated genes as perturbed genes. The purpose is to compare between differentially expressed genes (that are commonly considered important) and perturbed genes estimated by the three methods (DISCERN, LNS, and D-score), in terms of the enrichment for genes with potential importance to the disease. For ANOVA, in Fig. 3A-C, we considered 8,993 genes (AML), 7,922 genes (BRC) and 13,344 genes (LUAD) that show significant differential expression between cancer and normal samples at FDR corrected p-value < 0.05. The perturbed genes identified by DISCERN are more associated with survival than differentially expressed genes captured by ANOVA in AML and LUAD (Fig. 3).

In addition to the comparison with other methods – LNS and D-score – we also compare with frequently mutated genes and genes annotated to be involved in the respective cancer. We considered the following three gene sets: 1) a gene set constructed based on the gene-disease annotation database, Malacards [81], 2) genes known to have cancer-causing mutations based on the Cancer Gene Census [82], and 3) genes predicted to have driver mutations identified by MutSig [5] applied to The Cancer Genome Atlas (TCGA) data for the respective cancer type. The Malacards (gene set #1) and TCGA driver gene sets (#3) are generated for each cancer type – AML, breast cancer, or lung cancer. For example, for Malacards, we used the genes that are annotated to be involved in AML in Malacards to compare it with DISCERN genes identified in AML. Similarly, for the TCGA driver gene sets (#3), we used the AML TCGA data to identify the frequently mutated genes that are likely driver genes, and compared with high DISCERN-scoring genes in AML. We used the breast cancer TCGA data for BRC, and lung cancer TCGA data for LUAD. The Cancer Gene Census (CGC) gene set is a rigorously defined set of genes with multiple sources of evidence that its genes are cancer drivers in a single or multiple cancers.

For each cancer type, we compared these three sets of genes with the perturbed genes identified by DISCERN – 1,351 (AML), 2,137 (BRC), and 3,836 (LUAD) genes with high DISCERN scores – on the basis of the significance of the enrichment for survival-associated genes. S2 Fig shows that the perturbed genes identified by DISCERN are more significantly enriched for survival-associated genes.

### Prognostic model based on high DISCERN-scoring genes

In this section, we evaluated the DISCERN score based on how well it identifies genes that are predictive of patient prognosis. Here, we test the possibility of using the network perturbed genes identified by DISCERN as prognostic markers. For the cancer types with at least three data sets (AML and BRC; see Table 1), we construct a survival time prediction model using the genes with significant DISCERN scores (AML: 1,351 genes, BRC: 2,137 genes) identified based on one data set (Data # 1: AML1 and BRC1) as described in the previous subsection. Then, we trained the prediction model using one of the other datasets (Data #2: AML2 and BRC2) not used for the computation of the DISCERN score. Finally, we tested the prediction accuracy on the third data set (Data #3: AML3 and BRC3).

We controlled for clinical covariates whose data are available – age in case of AML and age, grade and subtype in case of BRC – by adding them as unpenalized covariates into our elastic net Cox regression model. We trained the Cox regression model using Data #2 and tested the survival prediction model on Data #3. Since we evaluated the survival prediction in separate data (AML3 and BRC3) that were not used in training the survival prediction model, using more predictors, e.g., by adding clinical covariates, does not necessarily improve the prediction performance. Adding more predictors often leads to a higher chance of overfitting. Our survival prediction model based on the high DISCERN-scoring genes works at least as well as models based on the genes contained in the previously established prognosis markers, such as Leukemic Stem Cell score (LSC) [54] for AML and MammaPrint signature (with ∼70 genes) [83] for BRC, as shown in Fig. 4. The c-index in AML is 0.669 with standard error (se) being 0.031 (Fig. 4B); in BRC, the c-index is 0.668 (se: 0.027) (Fig. 4D). The DISCERN-based expression marker with clinical covariates makes better predictions than when clinical covariates alone are used.

**Figure 4.**
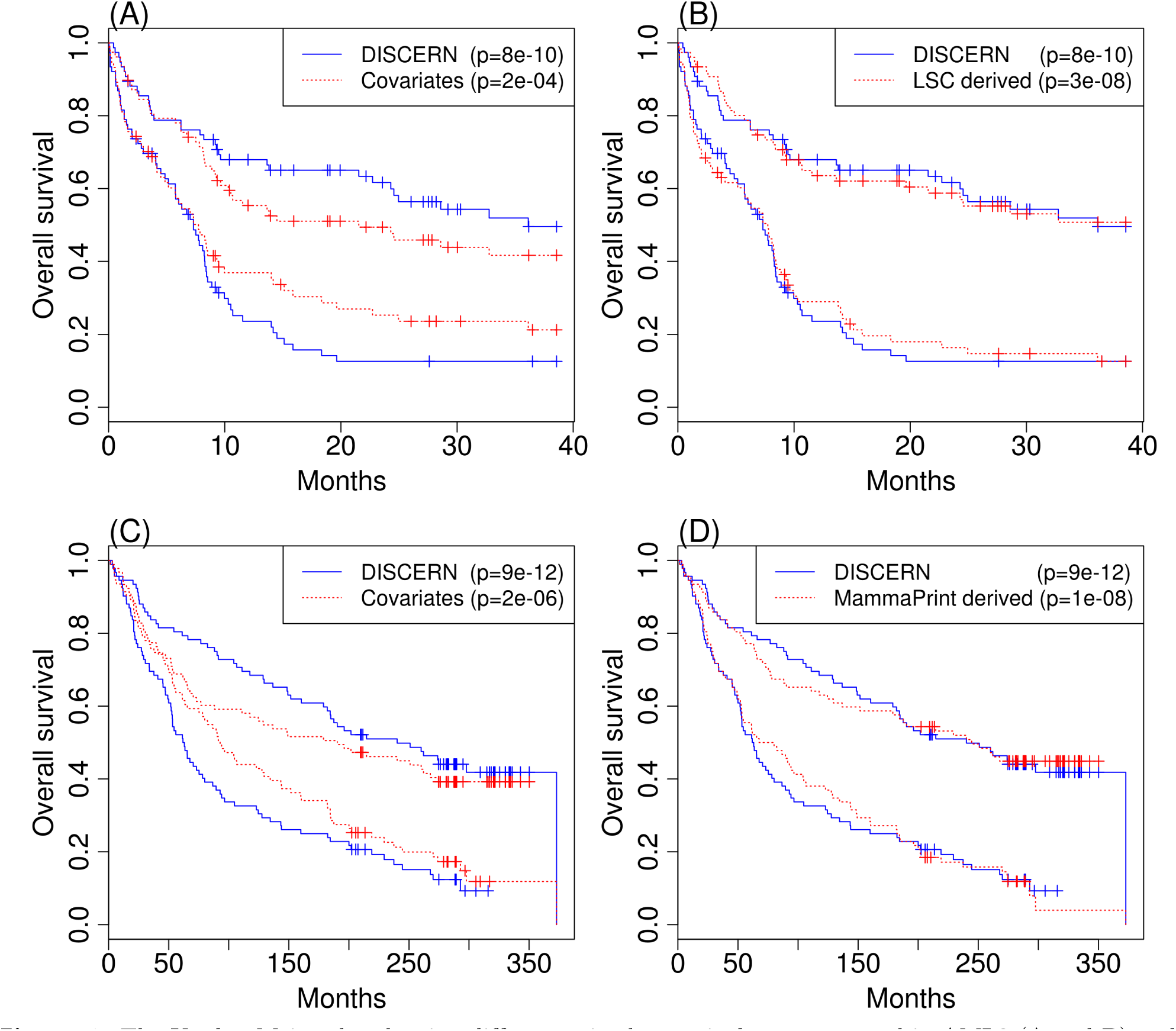
The Kaplan-Meier plot showing differences in the survival rate measured in AML3 (A and B) and BRC3 (C and D) between the two patient groups with equal size, created based on the predicted survival time from each prediction model. We consider the model trained based on the top N (=1,351 for AML; =2,137 for BRC) DISCERN-scoring genes and clinical covariates (blue), and the model trained based on only clinical covariates (red) (panels A and C for AML3 and BRC3, respectively). (B) The panel shows the comparison with the model trained using genes comprising 22 genes previously known prognostic marker, called LSC [54], along with the clinical covariates (red). (D) The panel shows the comparison with the model trained using 67 genes from the MammaPrint prognostic marker (70 genes) [83] along with the clinical covariates. We used 67 genes out of 70 genes that are present in our BRC expression datasets. P-values shown in each plot are based on the logrank test (red).

### DISCERN explains epigenomic activity patterns in cancer and normal cell types more accurately than alternative methods

One of the possible mechanisms underlying network perturbation identified in gene expression datasets representing different conditions (e.g., cancer and normal) is the following: A transcription factor (TF) ‘X’ binds to a gene ‘Y’s promoter or its enhancer region in cancer but not in normal (or vice versa). Then, ‘X’ or its co-regulator could be an expression regulator for ‘Y’ in cancer but not in normal (or vice versa), and Y would be identified as a perturbed gene (i.e., a high DISCERN-scoring genes). It is possible that ‘X”s binding information is not available and ‘X”s protein level is not reflected in its mRNA expression level; thus we cannot expect the DISCERN score of a gene inferred from expression data to be perfectly correlated with whether the gene has a differential biding of a certain TF, inferred from ChIP-seq or DNase-seq data. However, the degrees of correlation between the network perturbation score (DISCERN, LNS or D-score) of a gene and whether a TF differentially bind to the gene can be a way to evaluate the network perturbation scoring methods.

To determine whether or how much our statistical estimates of network perturbation reflect perturbation of the underlying TF regulatory network, we queried epigenomic data from ENCODE project. Two of the ENCODE cell lines – NB4 (an AML subtype [84]) and CD34+ (mobilized CD34 positive hematopoietic progenitor cells) – are closest to AML and normal conditions, and the DNase-seq data from these cell lines are available. We used the DNase-seq data from NB4 and the position weight matrices (PWMs) of 57 TFs available in the JASPAR database [85] to find the locations of the PWM motifs that are on the hypersensitive regions. This is a widely used approach to estimate active binding motifs using DNase-seq data, when ChIP-seq data are not available. We identified the locations of these PWM motifs on the hg38 assembly by using the FIMO [86] method (p-value ≤ 10^−5^). We then intersected these motif locations with hypersensitive regions identified by the DNase-seq data for each TF. We repeated for the other cell line CD34+.

For each TF, we measured how well the DISCERN score of a gene can predict the *differential* binding of the TF in active enhancer regions (marked by H3K27Ac) within 15kbs of the transcription start site (TSS) of the gene (Fig. 5A-C) and 5kb of the gene between blood cancer and normal cell lines (NB4 and CD34+) (**Fig. S3 FigA-C**). We show that the DISCERN score can reflect differential binding of most of the TFs better than existing methods to identify network perturbation (LNS and D-score) and a method to identify differentially expressed genes (ANOVA). As a way to summarize these results across all 57 TFs, we computed the Pearson’s correlation between the score of each gene and the proportion of TFs that differentially bind to that gene out of all TFs that bind to that gene. Fig. 5D shows that DISCERN detects genes with many TFs differentially bound between cancer and normal better than the other network perturbation detection methods (LNS and D-score) and ANOVA.

**Figure 5.**
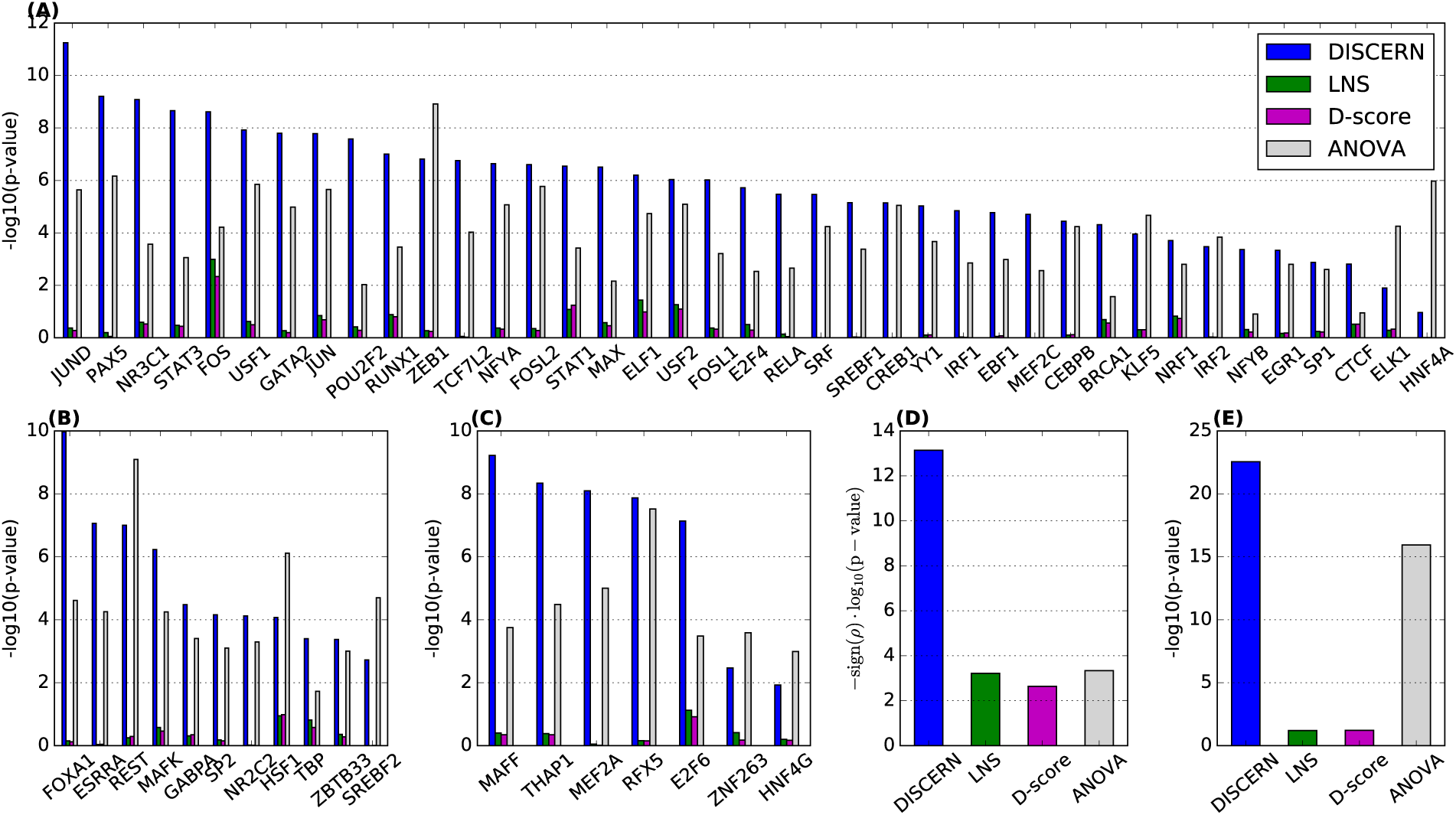
Kolmogorov-Smirnov test p-value measuring the significance of the difference in score between genes differentially bound by the corresponding transcription factor (TF) (x-axis) and those not differentially bound by the corresponding TF. We performed the one-sided test with an alternative hypothesis that differentially bound genes have higher scores; thus high - log_10_(p-value) means that high-scoring genes tend to show differential binding. The TFs are divided into the 3 sets: (A) TFs that are known to be associated with leukemia, (B) TFs that are known to be associated with cancer, and (C) TFs that are currently not known to be associated with cancer or leukemia, based on the gene-disease annotation database Malacards [81]. (D) Comparison of the p-values from the Pearson’s correlation between the score of each gene and the proportion of differential TFs out of all TFs bound to the genes. (E) Kolmogorov-Smirnov test (one-sided as above) p-value measuring the significance of the difference in score between the genes with differential binding purely based on the DNase-seq data and those not. Here, a differentially bound gene is defined as a gene that has a DNase signal within a 150bp window around its TSS in one condition but not in the other condition.

Considering hypersensitive sites identified by Dnase-seq data as the indication of “general” binding of TFs or other DNA-associated proteins, we assume that a gene is differentially bound if there is a DNase signal within a 150bp window around its TSS in one condition (cancer or normal), but not in the other condition. We observe that the DISCERN scores of the genes that are differentially bound are significantly higher than those of the genes that are not (Fig. 5E). These results suggest that DISCERN identifies possible regulatory mechanisms underlying network perturbation more accurately than existing network perturbation detection methods (LNS and D-Score) and a method for identifying differential expression levels (ANOVA).

As a specific example, STAT3 has been shown to differentially regulate the mRNA expression of BATF in myeloid leukemia but not in normal condition [87]. We found that STAT3 differentially binds to BATF in the AML cell line but not in the normal cell line based on our differential binding analysis using the DNase-seq/motif data, as described above (**S4 Table**). Interestingly, DISCERN identifies BATF as a perturbed gene in AML (FDR corrected p-value < 0.05). DISCERN also identifies STAT3 as the strongest regulator for BATF in AML expression data, but STAT3 is not selected as an expression regulator in normal expression data (Supporting Information). Interestingly, LNS and D-Score detect STAT3 as an expression regulator of BATF in both conditions, not as a differential expression regulator.

Two of the Tier 1 ENCODE cell lines – K562 (chronic myeloid leukemia cell line) and GM12878 (a lymphoblastoid cell line) – correspond to blood cancer and normal tissues as well [88]. Tier 1 data contains the largest number of TFs with ChIP-seq datasets, which allows us to perform this kind of analyses using ChIP-seq datasets for these TFs. We repeated the same analysis with these cell lines and showed similar results (see **S4 Fig** and **S5 Fig**).

### Combining DISCERN with ENCODE data improves the enrichment of known pathways

Additionally, we investigated whether one can use DISCERN as a filtering step to increase the power for a pathway enrichment analysis. We consider hypersensitive sites identified by DNase-seq data as the indication of “general” binding of TFs or other DNA-associated proteins, and important regulatory events. As describe above, we identified differentially regulated genes between AML and normal cell lines (NB4/ CD34+) by identifying gene that have DNase-seq peaks within 150bp around the TSS in one condition (cancer or normal), but not in the other condition. There are 3,394 differentially regulated genes selected based on the DNase-seq data, of which 339 are significant DISCERN genes (Supporting Information). Presumably, these disease specific targets should be enriched for pathways or categories that will help us understand mechanisms underlying the disease. Alternatively, some targets may be spurious, especially considering the use of cell lines that are not a perfect match to healthy and diseased bone marrow samples and experimental noise.

Here we attempt to identify differentially regulated genes between AML and normal samples, by integrating the information on the DNase-seq data (i.e., differentially bound genes) and significantly perturbed genes identified by DISCERN based on the expression datasets from AML samples and normal non-leukemic bone marrow samples. To show that combining these two pieces of information helps us to identify pathways that are specifically active in one condition not in the other, we compared the significance of the enrichment for Reactome pathways measured in fold enrichment between 1) 339 differentially bound DISCERN genes (intersection of 3,394 differentially bound genes and high DISCERN-scoring genes), and 2) 3,394 differentially bound genes. **S6 Fig** shows that for most of the pathways, using the intersection of differentially bound and perturbed genes increases the fold enrichment compared to when differentially bound genes were used (Wilcox p-value < 7 × 10^−5^).

Among the pathways, ‘platelet activation signalling and aggregation’ shows significant improvement in fold enrichment: 1) when differentially bound DISCERN genes were used (ƒ = 2.9; FDR q-value = 0.01), compared to 2) when differentially bound genes were used (ƒ = 1.03). It has been shown that the interactions between platelets and AML cells have considerable effects on metastasis, and the various platelet abnormalities have been observed in AML and other leukemias [89]. G-alpha signalling-related pathways also show significant boost in fold enrichment when DISCERN was used as a filtering mechanism for differentially bound genes. ‘G_q_ signalling pathway’ shows significant increase in fold enrichment: 1) when differentially bound DISCERN genes were used (ƒ = 2.16; FDR q-value = 0.05), compared to 2) when differentially bound genes were used (ƒ = 0.92). ‘G_12/13_ signalling pathway’ shows significant improvement in fold enrichment: 1) when differentially bound DISCERN genes were used (ƒ = 3.4; q-value < 0.03), compared to 2) when differentially bound genes were used (ƒ = 1.5). These pathways have been implicated in leukemias [90].

## Discussion

We presented a general computational framework for identifying the perturbed genes, i.e., genes whose network connections with other genes are significantly different across conditions, and tested the identified genes with statistical and biological benchmarks on multiple human cancers. Our method outperforms existing alternatives, such as LNS, D-score, and PLSNet, based on synthetic data experiments and through biological validation performed using seven distinct cancer genome-wide gene expression datasets, gathered on five different platforms and spanning three different cancer types – AML, breast cancer and lung cancer. We demonstrated that DISCERN is better than other methods for identifying network-perturbation in terms of identifying genes known to be or potentially important in cancer, as well as genes that are subject to differential binding of transcription factor according to the ENCODE DNase-seq data. We also demonstrated a method to use DISCERN scores to boost signal in the enrichment test of targets of differential regulation constructed using DNase-seq data available through the ENCODE Project.

## Methods

### Data Preprocessing

Raw cell intensity files (CEL) for gene expression data in AML1, AML2, and AML3 were retrieved from GEO [91] and The Cancer Genome Atlas (TCGA). Expression data was then processed using MAS5 normalization with the Affy Bioconductor package [92], and mapped to Enztrez gene annotations [93] using custom chip definition files (CDF) [94], and batch-effect corrected using ComBat [95] implemented in package `sva` from CRAN.

BRC1 was accessed through Broad Firehose pipeline (build 2013042100). We checked whether BRC1 processed by Firehose shows evidence of batch effects. We confirmed that the first three principal components are not significantly associated with the plate number (which we assumed to be a batch variable), which indicates no strong evidence of batch effects. BRC2 and BRC3 were accessed through Synapse (syn1688369, syn1688370). All probes were then filtered and mapped using the illuminaHumanv3.db Bioconductor package [96]. Probes mapped into the same genes were then collapsed by averaging if the probes being averaged were significantly correlated (Pearson’s correlation coefficient greater than 0.7).

LUAD1 was accessed through Broad Firehose pipeline (build 2015110100). Genes which had a very weak signal were filtered out of the LUAD1 data. We then applied the *voo*m normalization method that is specifically designed to adjust for the poor estimate of variance in count data, especially for genes with low counts [51]. The voom algorithm adjusts for this variance by estimating precision weights designed to adjust for the increased variance of observations of genes with low counts. This would stabilize the estimated distribution of RSEM values in the LUAD data, making it more normally distributed. Noting that LUAD data comes from different tissue source sites, we have applied batch-effect correction using ComBat.

For all datasets, only probes that map into genes that have Entrez gene names were considered. Table 1 shows the number of samples and genes used in each dataset. For AML1, BRC1, and LUAD1 that were used for score computationa, we splited each dataset into two matrices, one with only cancerous patients and one with normal patients. These matrices are normalized to 0-mean, unit-variance gene expression levels for each gene, before each network perturbation score (DISCERN, LNS, and D-score) was computed, which is a standard normalization step for accurately measuring the difference in the network connectivity. For methods that measure the differential expression levels (ANOVA), such normalization was not applied.

Lastly, candidate regulators are identified from a set of 3,545 genes known to be transcription factors, chromatin modifies, or perform other regulatory activity, which have been used in many studies on learning a gene network from high-dimensional expression data [43–46] **(S1 Table).**

### DISCERN score

DISCERN uses a likelihood-based scoring function that measures for each gene how much likely the gene is differently connected with other genes in the inferred network between two conditions (e.g., cancer and normal). We model each gene’s expression level based on the sparse linear model. Let 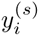 be a *standardized* expression levels of gene *i* in an individual with a condition *s* (cancer or normal) modeled as: 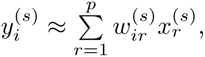 where 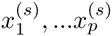 denote standardized expression levels of candidate regulator genes in a condition *s.* Standardization is a standard practice of normalizing expression levels of each gene to be mean zero and unit stadard deviation before applying penalized regression method [47–50]. To estimate weight vector 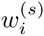 *lasso* [47] optimizes the following objective function: 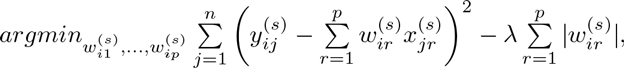 where the subscript *j* in the formula iterates over all patients, used as training instances for *lasso.* Here, 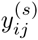 corresponds to the expression level of the *i*^*th*^ gene in the *j*^*th*^ patient in the *s*^*th*^ state and 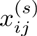 similarly corresponds to the expression level of the *i*^*th*^ regulator in the *j*^*th*^ patient in the *s*^*th*^ state. The second term, the L_1_ penalty function, will zero out many irrelevant regulators for a given gene, because it is known to induce sparsity in solution [47].

After estimating 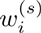 for each *s*, the DISCERN score measures how well each weight vector learned on one condition explains the data in the other condition, by using a novel model selection criteria defined as:

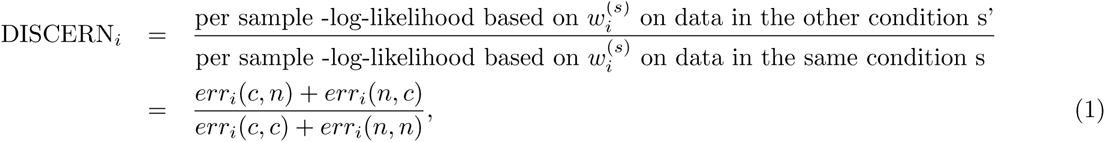

where 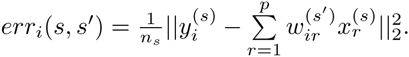. Here *n*_*s*_ is the number of samples in the data from condition *s*. The r=1 numerator in Eq (1) measures the error of predicting gene i’s expression levels in cancer (normal) based on the weights learned in normal (cancer). If gene i has different sets of regulators between cancer and normal, it would have a high DISCERN score. The denominator plays an important role as a normalization factor. To show that, we defined an alternative score, namely the *D*^0^ score that uses only the numerator of the DISCERN score, Eq (1):

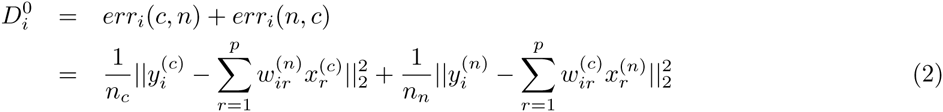

The first step of calculating the DISCERN score, and *D*^0^ score is to fit a sparse linear model (such as lasso [47]) for each gene’s expression level. We used the *scikit-learn* Python package (version 0.14.1) to calculate these scores with the values of the sparsity tuning parameters λ chosen by using the 5-fold cross-validation tests.

### ANOVA score computation

Analysis of Variance (ANOVA) is a standard statistical technique to measure the statistical significance of the difference in mean between two or more groups of numbers. For each gene, the 1-way ANOVA test produces a p-value from the F-test, which measures how significantly its expression level is different between conditions (e.g., cancer and normal). The ANOVA score was computed as negative logarithm of a p-value, obtained from 1-way ANOVA test using f oneway function in scipy.stats Python package.

### PLSNet score computation

PLSNet score attempts to measure how likely each gene is differently connected with other genes between conditions. It was computed using dna R package version 0.2_1 [34]. The network perturbation score for each gene is computed based on the empirical p-value from 1,000 permutation tests.

### LNS score computation

In Guan et al. (2013) [35], the authors defined the local network similarity (LNS) score for gene *i* that is defined as correlation of the Fisher’s z-transformed correlation coefficients between expression of gene *i* and all other genes between two conditions:

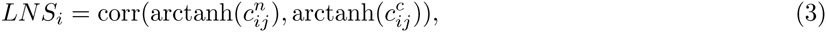

where 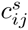 represents the correlation coefficient between expression levels of genes *i* and *j* in condition *s* = *n* for normal and *s* = *c* for cancer.

### D-score score computation

For synthetic data analysis, we have also introduced a D-score, computed as following (as used in Wang et al. (2009) [36]):

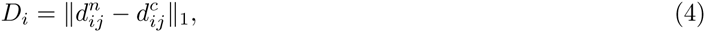

where 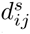 is a normalized correlation (normalized to have zero mean and unit variance across genes) between genes *i* and *j* in condition s, also known as Glass’ d score [97].

### Synthetic data generation

We generated 100 pairs of datasets, each representing disease and normal conditions. Each pair of datasets contains 100 variables drawn from the multivariate normal distribution with zero mean and covariance matrices Σ_1_ and Σ_2_. Each dataset contains *n*_1_ and *n*_2_, respectively, where *n*_1_ is randomly selected from uniform distribution between 100 and 110, and *n*_2_ is from uniform distribution between 16 and 26. This difference in *n*_1_ and *n*_2_ reflects the ratio of the cancer samples and normal samples in the gene expression data (Table 1).

For each of the 100 pairs of datasets, we divided 100 variables into the following three categories: 1) variables that have different sets of edge weights with other variables across two conditions, 2) variables that have exactly the same sets of edge weights with each other across the conditions, and 3) variables not connected with any other variables in the categories 2) and 3) in both conditions. For example, in Fig. 1A, ‘1’ is in category 1) (i.e., perturbed genes). ‘2’, ‘4’, ‘6’, and ‘7’ are in category 2), and ‘3’ and ‘5’ is in category 3). In each of the 100 pairs of datasets, the number of genes in category #1 (perturbed genes), *p*, is randomly selected from uniform distribution between 5 and 15. The number of genes in each of the other two categories #2 and #3 is determined as (100 - *p*)/2.

We describe below how we generated the network edge weights (i.e., elements of 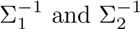) among the 100 variables. To ensure that only the genes in #1 have differing edge weights between two conditions, we generated two *p* × *p* matrices, *X*_1_ and *X*_2_, with elements randomly drawn from a uniform distribution between −1 and 1. Then, we generated symmetric matrices, 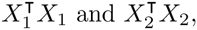, and added positive values to the diagonal elements to these symmetric matrices, if its minimum eigenvalue is negative – a commonly used method to generate positive definite matrices [98]. They become submatrices of 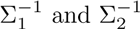 for these *p* variables. Similarly, we generate a common submatrix for the variables in category #2 – variables that have the same edge weights with other variables across conditions. Variables in category #3 have identify matrix as the inverse covariance matrix among the variables in that categories. Finally, we added mean zero Gaussian noise to each element of 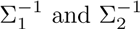 where the standard deviation of the Gaussian noise is randomly selected between 0.5 and 5.

This procedure allows having datasets of varying levels of difficulty in terms of high-dimensionality and network perturbation, which provides an opportunity to compare the average performances of the methods in various settings.

### Conservative permutation tests

To generate a conservative null distribution, we performed permutation tests by randomly reassigning cancer/normal labels to each sample, preserving the total numbers of cancer/normal samples. The correlation structure among genes would be preserved, because every gene is assigned the same permuted label in each permutation test. We then computed the DISCERN score for a random subset of 300 genes. We repeated this process to get over one million DISCERN scores to form a stable null distribution, which was used to compute empirical p-values.

### Identifying survival associated genes

For the survival-associated genes enrichment analysis, we first computed the association between survival time and each gene expression level. Genes that had a p-value from the Cox proportional hazards model (computed using *survival* R package) smaller than 0.01 were considered significantly associated with survival. These include 1,280 genes (AML), 1,891 genes (BRC) and 1,273 genes (LUAD) (S3 Table). Enrichment of overlap with top *N* DISCERN, LNS, D-score and ANOVA -scoring genes was computed by using the Fisher’s exact test based on the hypergeometric distribution function from scipy. stats Python package [99].

### Gene sets previously known to be important in cancer

We presented the results on the comparison with three sets of genes that are known to be important in cancer (**S2 Fig**). Here, we describe how we obtained these gene sets. First, Malacards genesets were constructed based on malacards.org website accessed in September 2012. Second, we used a set of 488 genes we downloaded from Catalogue of Somatic Mutations in Cancer website (CGC) [82]. For each cancer type, we considered the intersection between this list and the genes that are present in the expression data. Finally, a set of genes likely to contain driver mutations selected by MutSig was defined as those that pass q < 0.5 threshold based on 20141017 MutSig2.0 report from Broad Firehose.

### Cross-dataset survival time prediction

To evaluate the performance of the DISCERN score on identifying genes to be used in a prognosis prediction model, we trained the survival prediction model using one dataset and tested the model on an independent dataset (Fig. 4). To train the survival prediction model, we used the elastic net regression (α = 0.5) using glmnet CRAN package (version 1.9-8). Available clinical covariates – age for AML, and age, grade and subtype for BRC – were added as unpenalized covariates. Regularization parameter λ was chosen by using the built-in cross-validation function. Testing was always performed in the independent dataset with held-out samples from the dataset that was not used for training. For comparison, we trained the prediction model using 22 LSC genes [54] with age in AML, and 67 genes from the 70-gene-signature [83] (3 genes from the signature were missing in the dataset we were using) with clinical covariates (age, stage, and subtype) in BRC, as shown in Fig. 4B and Fig. 4D, respectively.

### Epigenomics analysis

The Encyclopedia of DNA Elements (ENCODE) is an international collaboration providing transcription factor binding and histone modification data in hundreds of different cell lines [100]. Data for ENCODE analysis were accessed through the UCSC Genome Browser data matrix [101] and processed using the BedTools and pybedtools packages [102,103]. Two of the ENCODE cell lines – NB4 (an AML subtype [84]) and CD34+ (mobilized CD34 positive hematopoietic progenitor cells) – are closest to AML and normal conditions, and the DNase-seq data from these cell lines are available.

For each cell line, we used the DNase-seq data and the position weight matrices (PWMs) of 57 transcription factors (TFs) available in the JASPAR database [85] to find the locations of the PWM motifs that are on the hypersensitive regions. We identified the locations of these PWM motifs on the hg38 assembly by using FIMO [86] (p-value ≤ 10^−5^). We then intersected these motif locations with hypersensitive regions identified by the DNase-seq data for each TF. We repeated this process to identify active binding motifs of the 57 TFs in each of the cell lines, NB4 and CD34+.

For each TF, we identified the genes the TF differentialy binds to between cancer and normal cell lines. We assumed that a certain TF is bound near a gene if the center of the peak is in the active enhancer regions (marked by H3K27Ac) within 15kbs of the transcription start site (TSS) of the gene or the 5kb around the gene’s transcription start site. We show that for most of the TFs, differentially bound genes have significantly high DISCERN scores than those not (Fig 5A-C).

The differential regulator score for each gene was computed by taking the number of differentially bound TFs and dividing it by the total number of TFs bound to the gene in any condition. We show that the differential regulator score is highly correlated with the DISCERN score (Fig 5D). For DNase-based analysis (Fig 5E), we defined a gene to be differentially regulated if hypersensitive sites detected by DNase-seq are within 150bp upstream of the gene in one condition and not in another.

### Reactome enrichment and DISCERN filtering

A set of 605 Reactome pathways was downloaded through Broad Molecular Signature Database (MSigDB) [59]. We postulate that hypersensitive sites identified by DNase-seq in a particular cell line indicate the regions where important regulatory events occur, such as transcription factor binding. We constructed the list of differentially regulated genes by comparing the hypersensitive sites identified by DNase-seq data between cancer and normal cell lines within 150bp upstream from TSS of each gene. For each pathway, we computed the fold enrichment 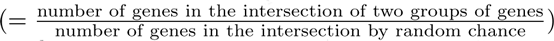 that measures the significance of the overlap between genes in the pathway and the identified differentially regulated genes. We compared the fold enrichment with when the genes in the intersection of differentially regulated genes and 1,351 significantly perturbed genes identified by DISCERN were used (**S6 Fig**). To reduce the noise, we only considered the pathways that had ≥ 5 genes in the overlap before filtering. The p-values were then FDR corrected for multiple hypothesis testing. Although p-values would measure the significance of the overlap between a gene set with a pathway, we used the enrichment fold as a measure of the significance of the overlap because we compared a set of genes with another set much smaller size.

## Acknowledgments

We would like to acknowledge Daniela Witten for her insightful advice on the development of the score function and Nao Hiranumi for sharing the locations data of JASPAR motifs he identified by running FIMO on the hg38 assembly.

This work was supported by a National Science Foundation (DBI-1355899), American Cancer Society (127332-RSG-15-097-01-TB), Solid Tumor Translational Research (Transformative Research Grant), and National Institutes of Health (U54CA149145).

## References

1. West J, Bianconi G, Severini S, Teschendorff AE. Differential network entropy reveals cancer system hallmarks. Scientific reports. 2012 Jan;2:802. Available from: www.nature.com/srep/2012/121113/srep00802/full/srep00802.html.

2. Schuster-Böckler B, Lehner B. Chromatin organization is a major influence on regional mutation rates in human cancer cells. Nature. 2012 Aug;488(7412):504–7. Available from: dx.doi.org/10.1038/nature11273.

3. Alexandrov LB, Nik-Zainal S, Wedge DC, Aparicio SAJR, Behjati S, Biankin AV, et al. Signatures of mutational processes in human cancer. Nature. 2013 Aug;Available from: www.nature.com/nature/journal/vaop/ncurrent/full/nature12477.html.

4. Baylin SB, Jones PA. A decade of exploring the cancer epigenome - biological and translational implications. Nature reviews Cancer. 2011 Oct;11(10):726–34. Available from: dx.doi.org/10.1038/nrc3130.

5. Lawrence MS, Stojanov P, Polak P, Kryukov GV, Cibulskis K, Sivachenko A, et al. Mutational heterogeneity in cancer and the search for new cancer-associated genes. Nature. 2013 Jul;499(7457):214–218. Letter. Available from: dx.doi.org/10.1038/nature12213.

6. Lovén J, Hoke HA, Lin CY, Lau A, Orlando DA, Vakoc CR, et al. Selective inhibition of tumor oncogenes by disruption of super-enhancers. Cell. 2013 Apr;153(2):320–34. Available from: www.cell.com/fulltext/S0092-8674(13)00393-0".

7. Stephens PJ, Tarpey PS, Davies H, Van Loo P, Greenman C, Wedge DC, et al. The landscape of cancer genes and mutational processes in breast cancer. Nature. 2012 Jun;486(7403):400–4.

8. Kim J, Kim I, Han SK, Bowie JU, Kim S. Network rewiring is an important mechanism of gene essentiality change. Scientific reports. 2012 Jan;2:900. Available from: www.nature.com/srep/2012/121129/srep00900/full/srep00900.html.

9. Califano A. Rewiring makes the difference. Molecular systems biology. 2011 Jan;7:463. Available from: dx.doi.org/10.1038/msb.2010.117.

10. Ideker T, Krogan NJ. Differential network biology. Molecular systems biology. 2012 Jan;8:565. Available from: dx.doi.org/10.1038/msb.2011.99.

11. Hanahan D, Weinberg RA. Hallmarks of cancer: the next generation. Cell. 2011 Mar;144(5):646–74. Available from: www.cell.com/fulltext/S0092-8674(11)00127-9.

12. Ciriello G, Cerami E, Sander C, Schultz N. Mutual exclusivity analysis identifies oncogenic network modules. Genome research. 2012 Feb;22(2):398–406. Available from: www.pubmedcentral.nih.gov/articlerender.fcgi?artid=3266046&tool=pmcentrez&rendertype=abstract.

13. Chapman MA, Lawrence MS, Keats JJ, Cibulskis K, Sougnez C, Schinzel AC, et al. Initial genome sequencing and analysis of multiple myeloma. Nature. 2011 Mar;471(7339):467–72. Available from: dx.doi.org/10.1038/nature09837.

14. Berger MF, Lawrence MS, Demichelis F, Drier Y, Cibulskis K, Sivachenko AY, et al. The genomic complexity of primary human prostate cancer. Nature. 2011 Feb;470(7333):214–20. Available from: dx.doi.org/10.1038/nature09744.

15. Bashashati A, Haffari G, Ding J, Ha G, Lui K, Rosner J, et al. DriverNet: uncovering the impact of somatic driver mutations on transcriptional networks in cancer. Genome biology. 2012 Dec;13(12):R124. Available from: genomebiology.com/2012/13/12/R124.

16. Ng S, Collisson EA, Sokolov A, Goldstein T, Gonzalez-Perez A, Lopez-Bigas N, et al. PARADIGM-SHIFT predicts the function of mutations in multiple cancers using pathway impact analysis. Bioinformatics (Oxford, England). 2012 Sep;28(18):i640–i646. Available from: www.pubmedcentral.nih.gov/articlerender.fcgi?artid=3436829&tool=pmcentrez&rendertype=abstract.

17. Vandin F, Upfal E, Raphael BJ. De novo discovery of mutated driver pathways in cancer. Genome research. 2012 Feb;22(2):375–85. Available from: genome.cshlp.org/content/22/2/375.long.

18. Carter H, Chen S, Isik L, Tyekucheva S, Velculescu VE, Kinzler KW, et al. Cancer-specific high-throughput annotation of somatic mutations: computational prediction of driver missense mutations. Cancer research. 2009 Aug;69(16):6660–7. Available from: cancerres.aacrjournals.org/content/69/16/6660.long.

19. Dees ND, Zhang Q, Kandoth C, Wendl MC, Schierding W, Koboldt DC, et al. MuSiC: identifying mutational significance in cancer genomes. Genome research. 2012 Aug;22(8):1589–98. Available from: www.pubmedcentral.nih.gov/articlerender.fcgi?artid=3409272&tool=pmcentrez&rendertype=abstract.

20. Stower H. Gene expression: Super enhancers. Nature reviews Genetics. 2013 Jun;14(6):367. Available from: www.nature.com.offcampus.lib.washington.edu/nrg/journal/v14/n6/full/nrg3496.html.

21. Whyte WA, Orlando DA, Hnisz D, Abraham BJ, Lin CY, Kagey MH, et al. Master transcription factors and mediator establish super-enhancers at key cell identity genes. Cell. 2013 Apr;153(2):307–19. Available from: www.cell.com/fulltext/S0092-8674(13)00392-9.

22. Won KJ, Zhang X, Wang T, Ding B, Raha D, Snyder M, et al. Comparative annotation of functional regions in the human genome using epigenomic data. Nucleic acids research. 2013 Apr;41(8):4423–32.

23. Zhu X, Ahmad SM, Aboukhalil A, Busser BW, Kim Y, Tansey TR, et al. Differential regulation of mesodermal gene expression by Drosophila cell type-specific Forkhead transcription factors. Development (Cambridge, England). 2012 Apr;139(8):1457–66. Available from: dev.biologists.org/content/139/8/1457.long.

24. Bhardwaj N, Kim PM, Gerstein MB. Rewiring of transcriptional regulatory networks: hierarchy, rather than connectivity, better reflects the importance of regulators. Science signaling. 2010 Jan;3(146):ra79. Available from: stke.sciencemag.org/cgi/content/abstract/3/146/ra79.

25. Shou C, Bhardwaj N, Lam HYK, Yan KK, Kim PM, Snyder M, et al. Measuring the evolutionary rewiring of biological networks. PLoS computational biology. 2011 Jan;7(1):e1001050. Available from: dx.plos.org/10.1371/journal.pcbi.1001050.

26. Habib N, Wapinski I, Margalit H, Regev A, Friedman N. A functional selection model explains evolutionary robustness despite plasticity in regulatory networks. Molecular systems biology. 2012 Jan;8:619. Available from: dx.doi.org/10.1038/msb.2012.50.

27. Przytycka TM, Singh M, Slonim DK. Toward the dynamic interactome: it’s about time. Briefings in bioinformatics. 2010 Jan;11(1):15–29.

28. Tusher VG, Tibshirani R, Chu G. Significance analysis of microarrays applied to the ionizing radiation response. Proceedings of the National Academy of Sciences of the United States of America. 2001 Apr;98(9):5116–21. Available from: www.pnas.org/content/98/9/5116.abstract.

29. Field A. Analysis of variance (ANOVA). Encyclopedia of measurement and statistics. 2007;p. 33–36.

30. Zhang BH, Liu J, Zhou QX, Zuo D, Wang Y. Analysis of differentially expressed genes in ductal carcinoma with DNA microarray. European review for medical and pharmacological sciences. 2013 Mar;17(6):758–66.

31. Mitra K, Carvunis AR, Ramesh SK, Ideker T. Integrative approaches for finding modular structure in biological networks. Nat Rev Genet. 2013 Oct;14(10):719–732.

32. Bockmayr M, Klauschen F, Györffy B, Denkert C, Budczies J. New network topology approaches reveal differential correlation patterns in breast cancer. BMC systems biology. 2013 Aug;7(1):78. Available from: www.biomedcentral.com/1752-0509/7/78.

33. Amar D, Safer H, Shamir R. Dissection of regulatory networks that are altered in disease via differential co-expression. PLoS computational biology. 2013 Jan;9(3):e1002955. Available from: dx.plos.org/10.1371/journal.pcbi.1002955.

34. Gill R, Datta S, Datta S. A statistical framework for differential network analysis from microarray data. BMC bioinformatics. 2010 Jan;11(1):95. Available from: biomedcentral.com/1471-2105/11/95.

35. Guan Y, Dunham MJ, Troyanskaya OG, Caudy AA. Comparative gene expression between two yeast species. BMC genomics. 2013 Jan;14:33. Available from: www.pubmedcentral.nih.gov/articlerender.fcgi?artid=3556494&tool=pmcentrez&rendertype=abstract.

36. Wang K, Narayanan M, Zhong H, Tompa M, Schadt EE, Zhu J. Meta-analysis of inter-species liver co-expression networks elucidates traits associated with common human diseases. PLoS computational biology. 2009 Dec;5(12):e1000616. Available from: dx.plos.org/10.1371/journal.pcbi.1000616.

37. Zhang B, Li H, Riggins RB, Zhan M, Xuan J, Zhang Z, et al. Differential dependency network analysis to identify condition-specific topological changes in biological networks. Bioinformatics (Oxford, England). 2009 Feb;25(4):526–32.

38. Li Y, Liang M, Zhang Z. Regression Analysis of Combined Gene Expression Regulation in Acute Myeloid Leukemia. PLoS Comput Biol. 2014 10;10(10):e1003908. Available from: dx.doi.org/10.1371%2Fjournal.pcbi.1003908.

39. Setty M, Helmy K, Khan AA, Silber J, Arvey A, Neezen F, et al. Inferring transcriptional and microRNA-mediated regulatory programs in glioblastoma. Mol Syst Biol. 2012 Aug;8:605–605. 22929615[pmid]. Available from: www.ncbi.nlm.nih.gov/pmc/articles/PMC3435504/.

40. Balwierz PJ, Pachkov M, Arnold P, Gruber AJ, Zavolan M, van Nimwegen E. ISMARA: automated modeling of genomic signals as a democracy of regulatory motifs. Genome Research. 2014;Available from: genome.cshlp.org/content/early/2014/03/25/gr.169508.113.abstract.

41. Meinshausen N, Buöhlmann P. High-dimensional graphs and variable selection with the Lasso. The Annals of Statistics. 2006 Jun;34(3):1436–1462.

42. Wainwright MJ, Lafferty JD, Ravikumar PK. High-Dimensional Graphical Model Selection Using L1-Regularized Logistic Regression. In: Advances in Neural Information Processing Systems; 2006.

43. Lee SI, Dudley AM, Drubin D, Silver PA, Krogan NJ, Pe’er D, et al. Learning a prior on regulatory potential from eQTL data. PLoS genetics. 2009 Jan;5(1):e1000358. Available from: dx.plos.org/10.1371/journal.pgen.1000358.

44. Jojic V, Shay T, Sylvia K, Zuk O, Sun X, Kang J, et al. Identification of transcriptional regulators in the mouse immune system. Nature immunology. 2013 Jun;14(6):633–43. Available from: dx.doi.org/10.1038/ni.2587.

45. Segal E, Shapira M, Regev A, Pe’er D, Botstein D, Koller D, et al. Module networks: identifying regulatory modules and their condition-specific regulators from gene expression data. Nature genetics. 2003 Jun;34(2):166–76. Available from: dx.doi.org/10.1038/ng1165.

46. Gentles AJ, Alizadeh AA, Lee SI, Myklebust JH, Shachaf CM, Shahbaba B, et al. A pluripotency signature predicts histologic transformation and influences survival in follicular lymphoma patients. Blood. 2009 Oct;114(15):3158–66. Available from: bloodjournal.hematologylibrary.org/content/114/15/3158.long.

47. Tibshirani R. Regression shrinkage and selection via the lasso. Journal of the Royal Statistical Society Series B (Methodological). 1996;p. 267–288.

48. Tibshirani R, Bien J, Friedman J, Hastie T, Simon N, Taylor J, et al. Strong rules for discarding predictors in lasso-type problems. Journal of the Royal Statistical Society: Series B (Statistical Methodology). 2012;74(2):245–266. Available from: dx.doi.org/10.1111/j.1467-9868.2011.01004.x.

49. Donald W Marquardt RDS. Ridge Regression in Practice. The American Statistician. 1975;29(1):3–20. Available from: www.jstor.org/stable/2683673.

50. Sardy S. On the Practice of Rescaling Covariates. International Statistical Review. 2008;76(2):285–297. Available from: dx.doi.org/10.1111/j.1751-5823.2008.00050.x.

51. Law CW, Chen Y, Shi W, Smyth GK. voom: Precision weights unlock linear model analysis tools for RNA-seq read counts. Genome Biol. 2014;15(2):R29.

52. Vogelstein B, Papadopoulos N, Velculescu VE, Zhou S, Diaz LA, Kinzler KW. Cancer genome landscapes. Science. 2013 Mar;339(6127):1546–1558.

53. Haferlach T, Kohlmann A, Wieczorek L, Basso G, Kronnie GT, Béné MC, et al. Clinical utility of microarray-based gene expression profiling in the diagnosis and subclassification of leukemia: report from the International Microarray Innovations in Leukemia Study Group. Journal of clinical oncology: official journal of the American Society of Clinical Oncology. 2010 May;28(15):2529–37. Available from: jco.ascopubs.org/content/28/15/2529.long.

54. Gentles AJ, Plevritis SK, Majeti R, Alizadeh AA. Association of a leukemic stem cell gene expression signature with clinical outcomes in acute myeloid leukemia. JAMA: the journal of the American Medical Association. 2010 Dec;304(24):2706–15. Available from: jama.jamanetwork.com/article.aspx?articleid=187113.

55. Gentles AJ, Plevritis SK, Majeti R, Alizadeh AA. Association of a leukemic stem cell gene expression signature with clinical outcomes in acute myeloid leukemia. JAMA. 2010 Dec;304(24):2706–2715.

56. Kharas MG, Lengner CJ, Al-Shahrour F, Bullinger L, Ball B, Zaidi S, et al. Musashi-2 regulates normal hematopoiesis and promotes aggressive myeloid leukemia. Nat Med. 2010 Aug;16(8):903–908.

57. Tanner SM, Austin JL, Leone G, Rush LJ, Plass C, Heinonen K, et al. BAALC, the human member of a novel mammalian neuroectoderm gene lineage, is implicated in hematopoiesis and acute leukemia. Proc Natl Acad Sci USA. 2001 Nov;98(24):13901–13906.

58. Alharbi RA, Pettengell R, Pandha HS, Morgan R. The role of HOX genes in normal hematopoiesis and acute leukemia. Leukemia. 2013 Apr;27(5):1000–1008.

59. Subramanian A, Tamayo P, Mootha VK, Mukherjee S, Ebert BL, Gillette MA, et al. Gene set enrichment analysis: a knowledge-based approach for interpreting genome-wide expression profiles. Proceedings of the National Academy of Sciences of the United States of America. 2005 Oct;102(43):15545–50. Available from: www.pnas.org/content/102/43/15545.abstract.

60. Jaatinen T, Laine J. Isolation of hematopoietic stem cells from human cord blood. Curr Protoc Stem Cell Biol. 2007 Jun;Chapter 2:Unit 2A.2.

61. Kottaridis PD, Gale RE, Frew ME, Harrison G, Langabeer SE, Belton AA, et al. The presence of a FLT3 internal tandem duplication in patients with acute myeloid leukemia (AML) adds important prognostic information to cytogenetic risk group and response to the first cycle of chemotherapy: analysis of 854 patients from the United Kingdom Medical Research Council AML 10 and 12 trials. Blood. 2001 Sep;98(6):1752–1759.

62. Karabon L, Pawlak E, Tomkiewicz A, Jedynak A, Passowicz-Muszynska E, Zajda K, et al. CTLA-4, CD28, and ICOS gene polymorphism associations with non-small-cell lung cancer. Hum Immunol. 2011 Oct;72(10):947–954.

63. Chen CH, Chuang SM, Yang MF, Liao JW, Yu SL, Chen JJ. A novel function of YWHAZ/Î^2^-catenin axis in promoting epithelial-mesenchymal transition and lung cancer metastasis. Mol Cancer Res. 2012 Oct;10(10):1319–1331.

64. Shiao YM, Chang YH, Liu YM, Li JC, Su JS, Liu KJ, et al. Dysregulation of GIMAP genes in non-small cell lung cancer. Lung Cancer. 2008 Dec;62(3):287–294.

65. Toyokawa G, Cho HS, Masuda K, Yamane Y, Yoshimatsu M, Hayami S, et al. Histone lysine methyltransferase Wolf-Hirschhorn syndrome candidate 1 is involved in human carcinogenesis through regulation of the Wnt pathway. Neoplasia (New York, NY). 2011 Oct;13(10):887–98.

66. Furukawa M, Soh J, Yamamoto H, Ichimura K, Shien K, Maki Y, et al. Silenced expression of NFKBIA in lung adenocarcinoma patients with a never-smoking history. Acta Med Okayama. 2013;67(1):19–24.

67. Xu HT, Wang L, Lin D, Liu Y, Liu N, Yuan XM, et al. Abnormal beta-catenin and reduced axin expression are associated with poor differentiation and progression in non-small cell lung cancer. Am J Clin Pathol. 2006 Apr;125(4):534–541.

68. Vire E, Brenner C, Deplus R, Blanchon L, Fraga M, Didelot C, et al. The Polycomb group protein EZH2 directly controls DNA methylation. Nature. 2006 Feb;439(7078):871–874.

69. Morin RD, Johnson NA, Severson TM, Mungall AJ, An J, Goya R, et al. Somatic mutations altering EZH2 (Tyr641) in follicular and diffuse large B-cell lymphomas of germinal-center origin. Nat Genet. 2010 Feb;42(2):181–185.

70. Kleer CG, Cao Q, Varambally S, Shen R, Ota I, Tomlins SA, et al. EZH2 is a marker of aggressive breast cancer and promotes neoplastic transformation of breast epithelial cells. Proc Natl Acad Sci USA. 2003 Sep;100(20):11606–11611.

71. Varambally S, Dhanasekaran SM, Zhou M, Barrette TR, Kumar-Sinha C, Sanda MG, et al. The polycomb group protein EZH2 is involved in progression of prostate cancer. Nature. 2002 Oct;419(6907):624–629.

72. Pujana MA, Han JD, Starita LM, Stevens KN, Tewari M, Ahn JS, et al. Network modeling links breast cancer susceptibility and centrosome dysfunction. Nat Genet. 2007 Nov;39(11):1338–1349.

73. Parker JS, Mullins M, Cheang MC, Leung S, Voduc D, Vickery T, et al. Supervised risk predictor of breast cancer based on intrinsic subtypes. J Clin Oncol. 2009 Mar;27(8):1160–1167.

74. van ‘t Veer LJ, Dai H, van de Vijver MJ, He YD, Hart AA, Mao M, et al. Gene expression profiling predicts clinical outcome of breast cancer. Nature. 2002 Jan;415(6871):530–536.

75. Gouge J, Satia K, Guthertz N, Widya M, Thompson AJ, Cousin P, et al. Redox Signaling by the RNA Polymerase III TFIIB-Related Factor Brf2. Cell. 2015 Dec;163(6):1375–1387.

76. Lagadec C, Meignan S, Adriaenssens E, Foveau B, Vanhecke E, Romon R, et al. TrkA overexpression enhances growth and metastasis of breast cancer cells. Oncogene. 2009 May;28(18):1960–1970.

77. Adriaenssens E, Vanhecke E, Saule P, Mougel A, Page A, Romon R, et al. Nerve growth factor is a potential therapeutic target in breast cancer. Cancer Res. 2008 Jan;68(2):346–351.

78. Paik S, Shak S, Tang G, Kim C, Baker J, Cronin M, et al. A Multigene Assay to Predict Recurrence of Tamoxifen-Treated, Node-Negative Breast Cancer. New England Journal of Medicine. 2004;351(27):2817–2826. PMID: 15591335. Available from: dx.doi.org/10.1056/NEJMoa041588.

79. van de Vijver MJ, He YD, van’t Veer LJ, Dai H, Hart AAM, Voskuil DW, et al. A gene-expression signature as a predictor of survival in breast cancer. The New England journal of medicine. 2002 Dec;347(25):1999–2009. Available from: www.ncbi.nlm.nih.gov/pubmed/12490681.

80. Therneau TM, Grambsch PM. Modeling Survival Data: Extending the Cox Model; 2000.

81. Rappaport N, Nativ N, Stelzer G, Twik M, Guan-Golan Y, Stein TI, et al. MalaCards: an integrated compendium for diseases and their annotation. Database: the journal of biological databases and curation. 2013 Jan;2013:bat018. Available from: www.pubmedcentral.nih.gov/articlerender.fcgi?artid=3625956&tool=pmcentrez&rendertype=abstract.

82. Futreal PA, Coin L, Marshall M, Down T, Hubbard T, Wooster R, et al. A census of human cancer genes. Nature reviews Cancer. 2004 Mar;4(3):177–83.

83. Glas AM, Floore A, Delahaye LJMJ, Witteveen AT, Pover RCF, Bakx N, et al. Converting a breast cancer microarray signature into a high-throughput diagnostic test. BMC genomics. 2006 Jan;7(1):278. Available from: www.biomedcentral.com/1471-2164/7/278.

84. Lanotte M, V MT, S N, P B, F V, R B. NB4, a maturation inducible cell line with t(15;17) marker isolated from a human acute promyelocytic leukemia (M3). Blood. 1991 Mar;77 (5):1080–6.

85. Sandelin A, Alkema W, Engström P, Wasserman WW, Lenhard B. JASPAR: an open-access database for eukaryotic transcription factor binding profiles. Nucleic Acids Research. 2004;32(suppl 1):D91–D94. Available from: nar.oxfordjournals.org/content/32/suppl_1/D91.abstract.

86. Grant CE, Bailey TL, Noble WS. FIMO: scanning for occurrences of a given motif. Bioinformatics. 2011;27(7):1017–1018. Available from: bioinformatics.oxfordjournals.org/content/27/7/1017.abstract.

87. Senga T, Iwamoto T, Humphrey SE, Yokota T, Taparowsky EJ, Hamaguchi M. Stat3-dependent induction of BATF in M1 mouse myeloid leukemia cells. Oncogene. 2002 Nov;21(53):8186–8191.

88. Thurman RE, Rynes E, Humbert R, Vierstra J, Maurano MT, Haugen E, et al. The accessible chromatin landscape of the human genome. Nature. 2012 Sep;489(7414):75–82.

89. Foss B, Bruserud O. Platelet functions and clinical effects in acute myelogenous leukemia. Thromb Haemost. 2008 Jan;99(1):27–37.

90. Siehler S. Regulation of RhoGEF proteins by G12/13-coupled receptors. Br J Pharmacol. 2009 Sep;158(1):41–49.

91. Barrett T, Wilhite SE, Ledoux P, Evangelista C, Kim IF, Tomashevsky M, et al. NCBI GEO: archive for functional genomics data sets–update. Nucleic acids research. 2013 Jan;41(Database issue):D991–5. Available from: nar.oxfordjournals.org/content/41/D1/D991.full.

92. Gautier L, Cope L, Bolstad BM, Irizarry RA. affy–analysis of Affymetrix GeneChip data at the probe level. Bioinformatics (Oxford, England). 2004 Feb;20(3):307–15.

93. Maglott D, Ostell J, Pruitt KD, Tatusova T. Entrez Gene: gene-centered information at NCBI. Nucleic acids research. 2011 Jan;39(Database issue):D52–7.

94. Dai M, Wang P, Boyd AD, Kostov G, Athey B, Jones EG, et al. Evolving gene/transcript definitions significantly alter the interpretation of GeneChip data. Nucleic acids research. 2005 Jan;33(20):e175.

95. Johnson WE, Li C, Rabinovic A. Adjusting batch effects in microarray expression data using empirical Bayes methods. Biostatistics (Oxford, England). 2007 Jan;8(1):118–27. Available from: biostatistics.oxfordjournals.org/content/8/1/118.long.

96. Dunning M, Lynch A, Eldridge M. illuminaHumanv3.db: Illumina HumanHT12v3 annotation data (chip illuminaHumanv3);.

97. Glass GV. Primary, secondary, and meta-analysis of research. Educational researcher. 1976;p. 3–8.

98. Boyd S, Vandenberghe L. Convex Optimization. New York, NY, USA: Cambridge University Press; 2004.

99. Jones E, Oliphant T, Peterson P, Others. {SciPy}: Open source scientific tools for {Python};.

100. ENCODE. The ENCODE (ENCyclopedia Of DNA Elements) Project. Science (New York, NY). 2004 Oct;306(5696):636–40.

101. Kent WJ, Sugnet CW, Furey TS, Roskin KM, Pringle TH, Zahler AM, et al. The Human Genome Browser at UCSC. Genome Research. 2002 May;12(6):996–1006.

102. Quinlan AR, Hall IM. BEDTools: a flexible suite of utilities for comparing genomic features. Bioinformatics (Oxford, England). 2010 Mar;26(6):841–2.

103. Dale RK, Pedersen BS, Quinlan AR. Pybedtools: a flexible Python library for manipulating genomic datasets and annotations. Bioinformatics (Oxford, England). 2011 Dec;27(24):3423–4.

